# Decoupling the bridge helix of Cas12a results in a reduced trimming activity and impaired conformational transitions

**DOI:** 10.1101/2021.02.25.432845

**Authors:** Elisabeth Wörle, Leonhard Jakob, Andreas Schmidbauer, Gabriel Zinner, Dina Grohmann

**Author notes:** For correspondence: Dina Grohmann, Department of Biochemistry, Genetics and Microbiology, Institute of Microbiology, University of Regensburg, Universitätsstraße 31, 93053 Regensburg, Germany.

## Abstract

The widespread and versatile prokaryotic CRISPR-Cas systems (clustered regularly interspaced short palindromic repeats and associated Cas proteins) constitute powerful weapons against foreign nucleic acids. Recently, the single-effector nuclease Cas12a that belongs to the type V CRISPR-Cas system was added to the Cas enzymes repertoire employed for gene editing purposes. Cas12a is a bilobal enzyme composed of the REC and Nuc lobe connected by a central structural element, the so-called bridge helix (BH). We generated BH mutants and integrated biochemical and single-molecule FRET (smFRET) studies to elucidate the role of the BH for the enzymatic activity and conformational flexibility of *Francisella novicida* Cas12a. We demonstrate that the BH impacts the trimming activity of Cas12a resulting in Cas12a variants with improved cleavage accuracy. Single-molecule FRET measurements reveal the hitherto unknown open and closed state of apo Cas12a. BH mutants preferentially adopt the open state. Transition to the closed state of the Cas12a-crRNA complex is inefficient in BH mutants but the semi-closed state of the ternary complex can be adopted even if the BH is deleted in its entirety. Taken together, these insights reveal that the BH is a structural element that influences the catalytic activity and impacts conformational transitions of FnCas12a.

## Introduction

Many bacteria and almost all archaea possess the adaptive immune system termed CRISPR-Cas (<u>c</u>lustered regularly interspaced <u>s</u>hort <u>p</u>alindromic repeats-<u>C</u>RISPR <u>as</u>sociated proteins) as part of their defense repertoire against viruses and mobile genetic elements (1–3). The CRISPR-Cas immune response is divided into three stages: I) Foreign nucleic acids (protospacers) are integrated as spacers between repeat sequences into the CRISPR-locus during the initial adaptation phase (4–6). II) In the expression and maturation phase, the CRISPR locus is transcribed and the resulting precursor CRISPR RNA (pre-crRNA) is further processed to yield the mature CRISPR RNA (crRNA)(7). III) In the final interference stage, the binary complex consisting of Cas protein(s) and crRNA binds invading DNA or RNA with sequence complementarity to the crRNA. This leads ultimately to the inactivation of the target nucleic acid most often as a result of endonucleolytic cleavage of the target DNA or RNA(1, 8).

The single effector nucleases Cas9, Cas12 and Cas13 garnered widespread attention as genome editing tools(9–13) because (i) Cas9 as well as Cas12a can be easily programmed to target a specific DNA sequence by complementary crRNAs and (ii) single effector nucleases act independently during the interference stage. Cas12a (formerly Cpf1) is an RNA-guided DNA endonuclease classified as class 2 type V-A Cas protein (3, 14, 15). Cas12a, in contrast to other Cas systems, is able to self-process the pre-crRNA (16). Programmed with a crRNA, Cas12a recognizes and binds a protospacer adjacent motif (PAM) in the target DNA with the sequence 5’-TTTV-3’ (V = G/C/A) (17, 18). The single active site for DNA cleavage resides in the RuvC domain, which cleaves the non-target strand (NTS) faster than the target strand (TS) resulting in a sequential cut thereby creating a double strand break with staggered ends (8, 19–22). After cleavage, Cas12a releases the PAM-distal part of the target DNA but remains bound to the PAM-proximal product (23, 24). This cleavage event is termed “cis-cleavage” of double-stranded DNA (dsDNA). However, Cas12a possesses also a trans-cleavage activity that is activated upon dsDNA cleavage. Here, a Cas12a-crRNA complex unspecifically binds to single-stranded DNA (ssDNA) and degrades it(25). The trans-cleavage activity of Cas12a has been repurposed to establish quantitative platforms for nucleic-acid detection (26–29).

Cas12a comprises a bilobal structure formed by the nuclease (Nuc) lobe (harboring the RuvC I, RuvC II, RuvC III, BH, and Nuc domains) and the recognition (REC) lobe (composed of the REC1 and REC2 domain). The wedge domain (WED I, WED II, WED III, PI (PAM interacting)) operates as hinge between the two lobes (Figure 1A) (2, 17, 30, 31). At the 5’ end, the crRNA forms a pseudoknot structure that is anchored in the WED domain and additionally forms an extensive network of interactions with the RuvC and REC2 domains (2, 30–32). The 3’-end of the crRNA encodes the sequence complementary to the target strand and interacts with the target DNA thereby forming an R-loop. Structural studies showed that Nuc and REC lobes undergo a closed to open transition upon loading of the DNA target in the crRNA-Cas12a complex (2, 17). The bridge helix (BH) is a central helix in the protein that structurally links the REC and the Nuc lobe. For Cas12a from *Acidaminococcus sp*. (AsCas12a) it has been shown that amino acids in the bridge helix interact with the sugar-phosphate backbone of the target DNA strand and play a role in the heteroduplex recognition of crRNA and target DNA(31). A tryptophan-residue at the end of the bridge helix (W958 in AsCas12a, W971 in Cas12a from *Francisella novicida* (FnCas12a)) is anchored in a hydrophobic pocket formed by the REC2 domain (Figure 1B/C). This interaction stabilizes the closed and active conformation of Cas12a (31, 32). The crRNA has a parallel orientation to the bridge helix and also connects the Nuc and the REC lobe (Figure 1A). In this context, the question remains if and how the BH influences the catalytic activity of Cas12a and whether the integrity of the BH as well as the anchoring in the REC lobe is required to ensure efficient cleavage activity of Cas12a. Furthermore, given the central position of the BH, the question arises whether the BH influences the conformational transitions of the enzyme.

**Figure 1.**
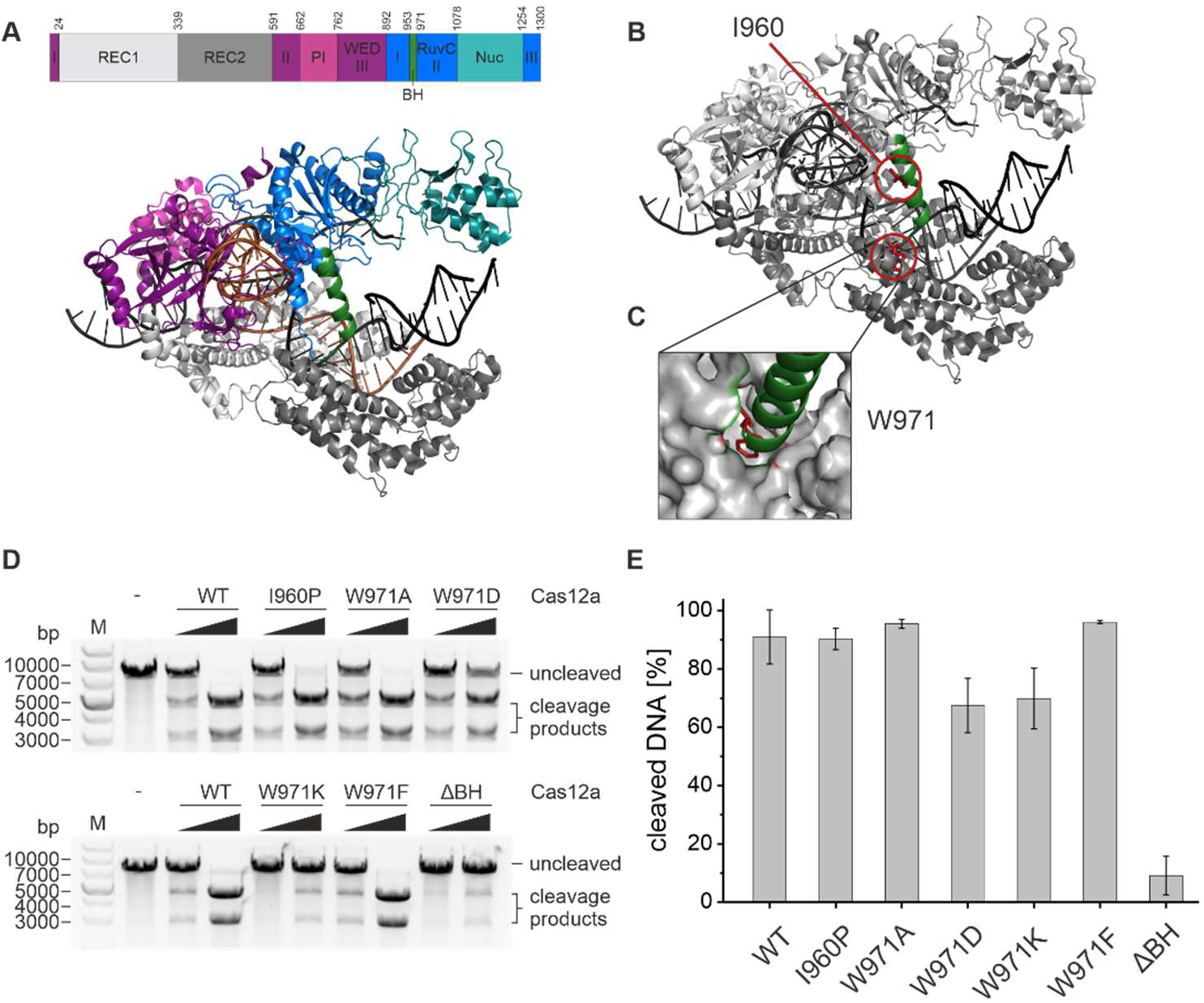
Structural organization of *Francisella novicida* Cas12a (FnCas12a) and influence of bridge helix mutations on Cas12a cleavage activity. **A** Domain organization and structure of FnCas12a in complex with crRNA (orange) and a double-stranded target DNA (black) (PDB: 6I1K). The bridge helix (BH) is highlighted in green. **B** Structure of Cas12a with residues I960 and W971 highlighted in red and the BH shown in green. **C** Close-up of the anchoring of W971 at the end of the bridge helix in a hydrophobic pocket in the REC2 domain. **D** Plasmid cleavage assay using WT Cas12a and Cas12a BH mutants using increasing concentrations of Cas12a (5 nM and 37.5 nM) and crRNA (5 nM and 37.5 nM) and target DNA at 5 nM. Incubation of the reaction at 37 °C for 1 h. The linearized target DNA plasmid (7300 bp) is cleaved into two products (4400 bp and 2900 bp). M, 1 kb plus DNA ladder; -, linearized target DNA without protein and crRNA. **E** Quantification of cleavage efficiency for the WT and mutant Cas12a variants for reactions containing 37.5 nM protein and crRNA. Shown is the average, error bars show the standard deviation of three independent assays.

Here, we performed a mutational analysis of the BH and evaluated the cleavage activity and cleavage accuracy of a variety of BH mutants. Cas12a is still catalytically active when the alpha-helical nature of the BH is disrupted or the anchoring of the BH in the REC lobe is prevented. However, mutations in the BH affect the trimming activity of We employed single-molecule Förster resonance energy transfer (smFRET) measurements on diffusing molecules to follow the conformational transitions of FnCas12a and of the BH mutants during their catalytical cycle making use of site-specifically labeled Cas12a. We show that Cas12a in its apo state exists in both, an open and closed conformation. The open conformation is preferred if the BH helix is not accurately anchored in the REC domain. Nevertheless, the crRNA induces the closure of the enzyme even if the anchoring of the BH is prevented or the BH is removed from the enzyme most likely because the adjacent helix 1 of the RuvC domain acts in concert with the BH and can ensure structural integrity. We demonstrate furthermore that the crRNA (independent of the BH) plays an essential role to direct the conformational transitions to reach a cleavage-competent state. A third, semi-closed conformation has to be adopted to allow the binding of target DNA. However, the stabilizing effect of the crRNA is sufficient to stabilize the semi-closed state in all BH mutants. Taken together, these data indicate that the BH is a structural element that influences the cleavage accuracy and trimming activity of Cas12a and enables the apo enzyme to adopt the closed state thereby promoting efficient crRNA loading.

## Material and Methods

### Protein expression and purification

The Cas12a gene from *Francisella novicida* was amplified from the plasmid pFnCpf1_min (Addgene #69975) (1) and cloned into pGEX-2TK via the BamHI and EcoRI restriction sites. This resulted in an expression plasmid that encoded the wildtype Cas12a sequence with an N-terminal GST-tag and Thrombin cleavage site. Cas12a mutants were (E1006A (dCas12a), I960P, W971A, W971D, W971K, W971F, ΔY953-K969 (ΔBH), ΔY953-W971 (ΔΔBH)) generated via site-directed mutagenesis (Table S1). Protein expression was performed in BL21(DE3) cells. The cells were grown at 37 °C. At an optical density (600 nm) of 0.4, the cells were transferred to 18 °C. Protein expression was induced with IPTG (1 mg/l) at an OD_600_ = 0.6. Cells were further incubated at 18 °C overnight. After harvesting the cells (10 min, 6000 g, 4 °C), they were resuspended in 20 ml Tris-Cas-A buffer (20 mM Tris/HCl pH 7.5, 50 mM NaCl, 5 mM MgCl_2_, 5% (v/v) Glycerol). 3 µl Universal Nuclease Mix (Pierce^™^) and 1 tablet of cOmplete^™^ Protease Inhibitor Cocktail (Roche) were added and the cells were lysed via sonication. The lysate was centrifuged (1 h, 30 000 g, 4 °C) and the supernatant applied to a GSTrap 5 ml FF column affinity purification column (GE Healthcare) at 4 °C equilibrated with Tris-Cas-A buffer. After binding of the protein, the column was washed with a high salt washing buffer (Tris-Cas-A buffer containing 1 M NaCl) and eluted with 100% Tris-Cas-B buffer (20 mM Tris-HCl pH 7.5, 50 mM NaCl, 5 mM MgCl_2_, 5% (v/v) Glycerol, 10 mM Glutathione). The eluted protein was incubated overnight with Thrombin protease (100 U/ml, 50:1 (v:v)) at 4 °C to allow the cleavage of the GST-tag. The GST-tag and Thrombin protease were removed via heparin affinity chromatography (HiTrap Heparin 5 ml FF, GE Healthcare). Cas12a was eluted via an elution gradient with 1 M NaCl (20 mM Tris/HCl pH 7.5, 1 M NaCl, 5 mM MgCl_2_, 5% (v/v) Glycerol). Cas12a typically eluted at a NaCl concentration of 760 mM. Protein concentrations were calculated using an extinction coefficient at 280 nm of 143 830 M^-1^cm^-1^ (ProtParam tool from www.expasy.org). Protein aliquots were flash frozen and the protein stored at −80 °C until further use.

Site-specific incorporation of the unnatural amino acid *para*-azido-L-phenylalanine into Cas12a To produce fluorescently labeled protein variants of Cas12a for single-molecule FRET measurements, the Amber-suppressor strategy was employed to site-specifically incorporate the unnatural amino acid *para*-azido-L-phenylalanine (AzF) (39, 40). The amber-stop-codon (TAG) was implemented either into the wildtype pGEX-2TK-Cas12a plasmid or the Cas12a plasmids already carrying mutations in the bridge helix via site-directed mutagenesis at positions D470, K647, and/or T1222 (Table S1). The plasmids with amber-codons were expressed in BL21(DE3), which additionally harbored the pEVOL-pAzF plasmid (Addgene #31186, (55)). This plasmid encodes an additional arabinose-inducible promoter that governs the expression of an amber-suppressor tRNA (tRNA_CUA_) and a biorthogonal tRNA synthetase. Expression and protein purification were carried out as described before with the following modification: additionally, 200 mg/l AzF were added to the expression culture at OD_600_ = 0.3 and L-Arabinose at a final concentration of 20 mg/l at OD_600_ = 0.4. Aliquots of the purified protein were flash frozen and stored at −80 °C.

### Protein labeling via unnatural amino acids incorporated into Cas12a

Cas12a variants with AzF at positions D470, K647, and/or T1222 were stochastically labeled via the Staudinger-Bertozzi ligation between a triphenylphosphine (at the DyLight fluorophores) and an azide moiety at AzF (41). The purified protein was incubated with a final concentration of 50 µM of DyLight 550 and DyLight 650 for 90 min at room temperature in darkness. The labelling reaction was centrifuged for 10 min at 21 000 g. The supernatant was purified via size exclusion chromatography using a Sephadex G-50 illustra NICK column and Cas batch buffer (50 mM Tris/HCl pH 7.5, 200 mM NaCl, 5 mM MgCl_2_, 2% (v/v) Glycerol, 0.1% (v/v) Tween20) to remove the excess of dye. The labeled protein was directly used for single molecule measurements.

### Synthetic oligonucleotides

Oligodeoxyribonucleotides were purchased from Eurofins-MWG (Ebersberg, Germany). The target DNA sequence was derived from Swarts and Jinek, 2018 (TS 5’-ACTCAATTTTGACAGCCCACATGGCATTCCACTTATCACTAAAGGCATCCTTCCACGT-3’, NTS 5’-ACGTGGAAGGATGCCTTTAGTGATAAGTGGAATGCCATGTGGGCTGTCAAAATTGAGT-3’) (21). For cleavage assays, the target strand carried a Cy5 label and the non-target strand a Cy3 label either on the 5’ end or the 3’ end. Target and non-target strand were annealed by incubation in annealing buffer (final concentration: 3 mM HEPES pH 7.4, 10 mM Potassium Acetate, 0.2 mM Magnesium Acetate) at 95 °C for 3 min and passive cool-down to room temperature.

### In vitro transcription

crRNAs (crRNA 5’-UAAUUUCUACUGUUGUAGAUGUGAUAAGUGGAAUGCCAUGUGGG-3’, pre-crRNA 5’-GCUGAUUUAGGCAAAAACGGGUCUAAGAACUUUAAAUAAUUUCUACUGUUGUAGAUGUGAUAAGUGGAAUGCCAUGUGGG-3’ derived from Creutzburg *et al*. (56)) were prepared and purified using the T7 RiboMAX^™^ Express Large Scale RNA Production System (Promega). The DNA templates (T7crRNA 5’-CCCACATGGCATTCCACTTATCACATCTACAACAGTAGAAATTACCCTATAGTGAGTC GTATTATCGATC-3’, T7pre-crRNA 5’-CCCACATGGCATTCCACTTATCACATCTACAACAGTAGAAATTATTTAAAGTTCTTAGACCCGTTTTTGCCTAAATCAGCCCCTATAGTGAGTCGTATTATCGATC-3’) were purchased from Eurofins-MWG (Ebersberg, Germany). The concentration was photometrically determined and the RNA stored at −20 °C.

### Electrophoretic mobility shift assay

The binary complex composed of 200 nM Cas12a protein and 200 nM crRNA was pre-formed in 1x Cas buffer (20 mM Tris/HCl pH 7.5, 100 mM NaCl, 10 mM MgCl_2_, 2% (v/v) Glycerin, 1 mM DTT, 0.05% (v/v) Tween20) for 10 min at room temperature. 75 nM of the binary complex were added to 10 nM of fluorescently labeled short target DNA in a total volume of 10 µl and incubated at 37 °C for 1 h. To avoid unspecific interactions of Cas12a with nucleic acids, Heparin with a final concentration of 32 µg/ml was added and the reaction incubated for additional 10 min at room temperature. Gel loading dye (final concentration: 62.5 mM Tris/HCl pH 6.8, 2.5% (w/v) Ficoll400) was added and the samples separated by a non-denaturing Tris-Glycine gel electrophoresis (6% PAA, 230 V, 30 min). The gel was visualized using a ChemiDoc Imaging System (Bio-Rad).

### Plasmid cleavage assay

To perform plasmid cleavage assays, the binary complex composed of Cas12a and crRNA was pre-formed by incubating 100 nM of protein and 100 nM crRNA for 10 min at room temperature in 1x Cas buffer (20 mM Tris/HCl pH 7.5, 100 mM NaCl, 10 mM MgCl_2_, 2% (v/v) Glycerin, 1 mM DTT, 0.05% (v/v) Tween20). 5 nM or 37.5 nM of preincubated binary complexes were added to 5 nM of target DNA (eGFP-hAgo2 plasmid (Addgene #21981) linearized with SmaI). The reaction was incubated in 1x Cas buffer in a total volume of 15 µl for 1 h at 37 °C. The reaction was stopped with 0.36 U Proteinase K. After incubation for 30 min at 55 °C, the reaction was mixed with loading dye, and loaded onto a 0.8% agarose 1x TAE gel (100 V, 30 min).

In order to follow cleavage kinetics, 25 nM of the binary complex were added to 5 nM of target DNA and incubated at 28 °C. After 15 s, 30 s, 1 min, 2 min, 3 min, 4 min, 5 min, 10 min, 15 min, 30 min, 1 h, and 2 h aliquots of the reaction were stopped with EDTA (final concentration 83.3 mM) and 0.36 U Proteinase K. After incubation for 30 min at 55 °C, the reaction was mixed with loading dye, and loaded onto a 0.8% agarose 1x TAE gel (100 V, 30 min).

### Short DNA target cleavage assay

The binary complex of 200 nM Cas12a protein and 200 nM crRNA was pre-formed in 1x Cas buffer for 10 min at room temperature. In a final concentration of 75 nM, the binary complex was added to 10 nM fluorescently labeled double-stranded short target DNA in a total volume of 10 µl and incubated at 37 °C for 1 h. The reaction was stopped by the addition of 0.36 U Proteinase K and incubated at 55 °C for 30 min. Loading dye (final concentration: 47.5% (v/v) formamide, 0.01% (w/v) SDS, 0.01% (w/v) bromophenol blue, 0.005% (w/v) xylene cyanol, 0.5 mM EDTA) was added and the samples were separated on a pre-heated 15% PAA, 6 M Urea, 1x TBE gel (300 V, 30 min). Fluorescent signals were visualized using a ChemiDoc Imaging System (Bio-Rad).

In order to follow cleavage kinetics, reactions were stopped after 1 s, 30 s, 1 min, 3 min, 5 min, 10 min, 20 min, 30 min, 45 min, and 60 min with EDTA (final concentration 125 mM) and 0.36 U Proteinase K. Further steps were conducted according to the cleavage assay as described above.

The cleavage reactions for the high-resolution sequencing gel (15% PAA, 6 M Urea, 1x TBE) were carried out with 375 nM of the binary complex and 50 nM of fluorescently labeled target DNA. After 1 h at 37 °C the reaction was stopped by the addition of 0.36 U Proteinase K and incubated at 55 °C for 30 min. Loading dye (final concentration: 47.5% (v/v) formamide, 0.01% (w/v) SDS, 0.01% (w/v) bromophenol blue, 0.005% (w/v) xylene cyanol, 0.5 mM EDTA) was added and the samples were separated on the pre-heated sequencing gel (45 °C, 65 W, 1 h). Fluorescent signals were visualized using a ChemiDoc Imaging System (Bio-Rad).

### Protein Thermal Shift^™^ melting curves

Melting curves of Cas12a variants were generated using the Protein Thermal Shift^™^ kit (ThermoFisher) using the Rotor-Gene Q qPCR cycler from Qiagen and the HRM (High Resolution Melt) software. The proteins were diluted in water and measurements were performed in quadruplicates. The data were analyzed using the Origin software.

### Confocal single-molecule FRET measurements

Sample chambers for confocal measurements (Cellview slide, Greiner Bio-One) were passivated with a passivation solution (2 mg/ml BSA in PBS) for 10 min and subsequently washed with 1x Cas buffer (20 mM Tris/HCl pH 7.5, 100 mM NaCl, 10 mM MgCl_2_, 2% (v/v) Glycerin, 1 mM DTT, 0.05% (v/v) Tween20). Labelled Cas12a apo protein (pM-nM) was diluted to picomolar concentrations in 1x Cas buffer and transferred into the sample chamber. The binary complex composed of Cas12a protein with crRNA (final concentration 0.5 nM, 1 nM, 2 nM, or 4 nM) was pre-incubated for 30 min at room temperature in a small volume of 20 µl and prior to the measurement diluted by a factor of 10 with Cas buffer to yield a final volume of 200 µl. For the ternary complexes, the binary complex composed of Cas12a with crRNA (final concentration 1 nM) was pre-incubated for 10 min prior to addition of double-stranded or single-stranded target DNA (final concentration 1 nM, 2 nM, or 4 nM) and incubated in a volume of 20 µl for 20 min. Prior to the measurements, the sample was diluted by a factor of 10 with 1x Cas buffer to yield the final volume of 200 µl. Single-molecule FRET measurements of freely diffusing molecules were carried out for 30 min.

The single-molecule measurements were conducted on a MicroTime 200 confocal microscope (PicoQuant) with pulsed laser diodes (532 nm: LDH-P-FA-530B; 636 nm: LDH-D-C-640; PicoQuant / clean-up filter: ZET 635; Chroma). Excitation of the fluorophores was done via pulsed interleaved excitation (PIE) with a laser intensity of 20 µW. To collect the emitted fluorescence, a 1.2 NA, x60 microscope objective (UplanSApo x60/1.2W; Olympus) and a 50 µm confocal pinhole were used. The donor and acceptor fluorescence were separated by a dichroic mirror (T635lpxr; Chroma) and bandpass filters (donor: ff01-582/64; Chroma; acceptor: H690/70; Chroma). An individual APD (SPCM-AQRH-14-TR, Excelitas Technologies) for each photon stream detected the separated photon streams, which were subsequently analyzed by a TCSPC-capable PicoQuant HydraHarp 400.

### Data analysis

Confocal smFRET data of diffusing molecules were analyzed with the software package PAM (57). Photon bursts were extracted by an all-photon burst search (APBS, parameters: *L* = 100, *M* = 15, *T* = 500 µs) and a dual-channel burst search (DCBS, parameters: *L* = 100, *M*_*GG+GR*_ = 15, *M*_*RR*_ = 15, *T* = 500 µs) (58). The obtained burst data were corrected for donor leakage and acceptor direct excitation, determined from the APBS data (59). β and γ factors were determined by applying an internal fit on distinct FRET populations in the ES-histograms obtained by the DCBS algorithm. The corrected DCBS data were binned (bin size = 0.032) and the mean value of triplicate measurements plotted as an E-histogram. The histograms were fitted with either one, two, or three Gaussian fits using the Origin software.

## Results

### Mutations in the bridge helix of Cas12a alter the trimming activity and cleavage rate

In order to shed light on the function and importance of the bridge helix (BH), a series of mutations were introduced in FnCas12a (Figure 1B). The I960P mutant was generated to disrupt the α-helical nature of the BH. The tryptophan at position 971 is accommodated in the hydrophobic pocket of the REC2 domain (31, 32) (Figure 1C). This residue was mutated to a range of amino acids with different chemical properties (W971A, W971D, W971K, W971F), e.g. amino acids that differ in size and charge, to investigate whether the anchoring of the bridge helix is important for the structural integrity and catalytic function of Cas12a. Additionally, we created a mutant in which the BH is deleted (ΔBH, ΔY953-K969, deletion of 17 amino acids). All mutants could be purified to homogeneity (Figure S1) and exhibited comparable stability as assessed by a thermal shift assay (Figure S2).

The activity of wildtype (WT) Cas12a and the BH mutants was analyzed in plasmid cleavage assays utilizing linearized plasmid DNA as target (Figure 1D, Figure S3). Addition of WT Cas12a resulted in the expected formation of two cleavage products (Figure 1D). All BH mutants were catalytically active but showed clear differences in cleavage efficiency. The ΔBH mutant was the least active mutant with only 15% of target DNA being cleaved as compared to 91% cleaved by the WT enzyme after 60 minutes (Figure 1E). EMSAs revealed that the ΔBH mutant is not impaired in DNA binding (Figure S4) suggesting that the reduced cleavage activity is solely caused by a catalytic defect of the mutants. Mutant W971A and W971F are not impaired in their activity. The I960P mutant was only marginally less active than the WT. Mutants W971D and W971K remained active and showed approximately 70% of the WT activity. A detailed study of the reaction kinetics was performed at 28 °C to be able to follow the reaction with a good time resolution (Figure S3). Here, cleavage products appeared already after 2 min for the WT enzyme. Variants W971K, W971D as well as I960P cleaved the DNA considerably slower than the WT. Interestingly, mutant W971F cleaved the target slightly faster than the wildtype and 80% of the substrate was cleaved (WT: 45%) 2h of incubation. The ΔBH was not catalytically active under these conditions.

In order to evaluate the cleavage pattern created by the BH mutants, we made use of a cleavage assay established by Swarts *et al*. based on fluorescently labeled short DNA target strands (58 nt, Figure 2A) (21). To follow the cleavage reaction on both strands simultaneously, a Cy3 and Cy5 label was attached to the 5’ end of the non-template strand (NTS) and the template strand (TS), respectively. Cleavage products were resolved on a denaturing sequencing gel in order to analyze cleavage products with single-nucleotide resolution (Figure 2B, Figure S5 and S6). In case of the WT enzyme, a predominant cleavage product is visible and two minor products with one and two additional nucleotides are detected. In contrast, the mutants show mainly longer cleavage products (up to 5 nt longer than the main product produced by the WT). Only the W971A and W971F mutants also generate products as short as the WT. Notably, this is not the main product of these mutants. Kinetic studies using the WT enzyme (Figure S5C/E) showed that also the WT produces longer NTS cleavage products at the start of the reaction. Over time, the long cleavage products are shortened to yield the main product. This is the result of a trimming activity that shortens the NTS by up to 4 nt (19). After cleavage of the target DNA, the PAM distal part of the DNA is released but Cas12a remains bound to the PAM proximal part (24) (here labeled with Cy3 at the NTS). While still bound, Cas12a further shortens the NTS via the trimming activity. Trimming of the NTS was also observed for the W971A and W971F mutants yielding two or three main cleavage products including the final cleavage product observed for the WT (Figure S6). However, trimming of the NTS by the mutated enzymes is less efficient as the final product is not the dominant product even after prolonged reaction times. In contrast, all other mutants never yielded the final cleavage product of the WT enzyme. This suggests that mutants I960P, W971D, W971K and ΔBH lack the trimming activity altogether. The cleavage pattern for the TS is the same for all Cas12a variants and shows one main cleavage product and two additional products 1 nt longer and 1 nt shorter than the main product. Our data are in agreement with previous studies that reported the imprecise cleavage of the TS by AsCas12a and FnCas12a (19, 20).

**Figure 2.**
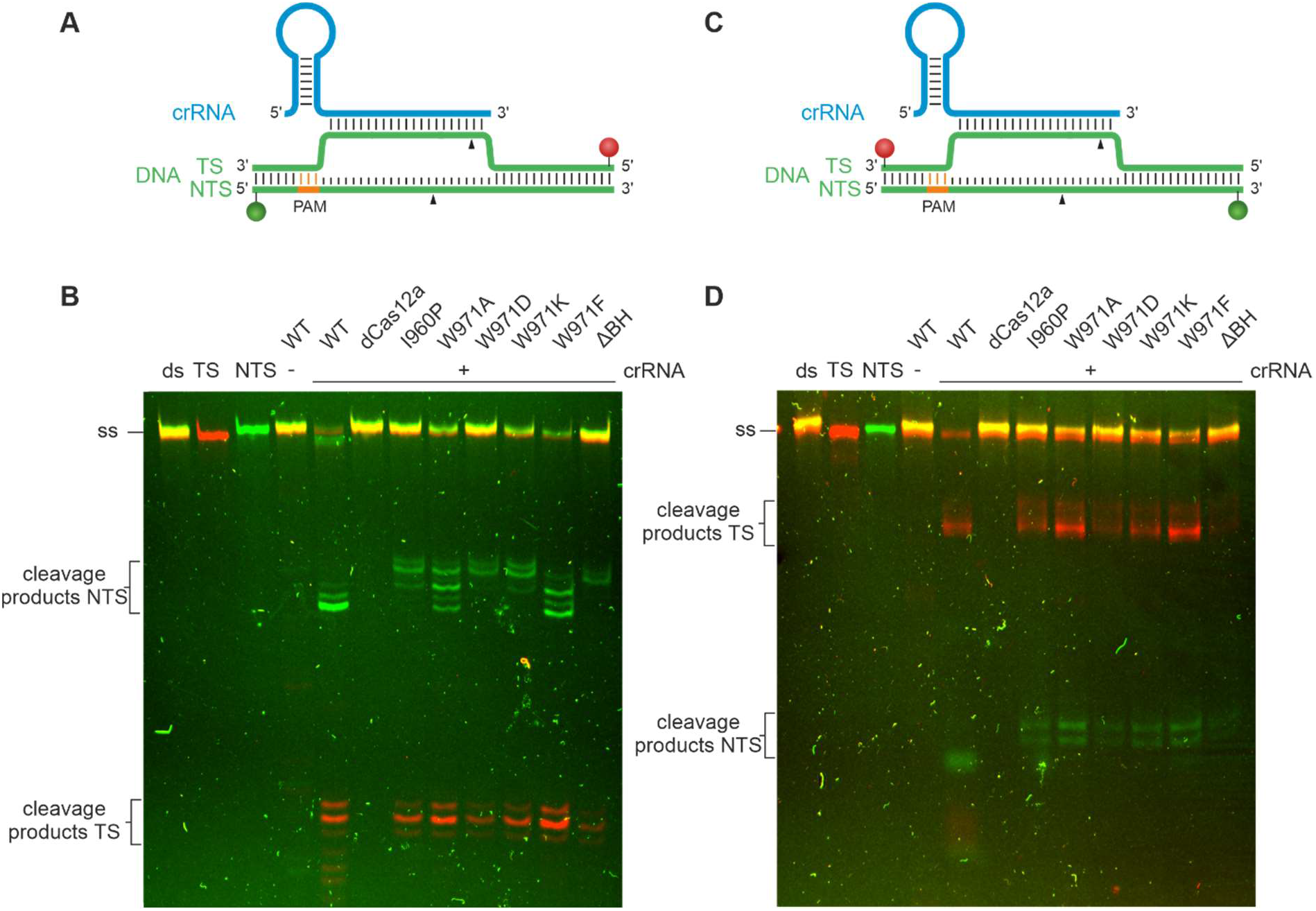
DNA cleavage assay reports on cleavage accuracy and trimming activity of Cas12a variants. Cleavage assay of WT Cas12a and Cas12a bridge helix mutants using a 7.5-fold excess of protein and crRNA (750 nM) relative to the target DNA (10 nM). The reaction was incubated at 37 °C for 1 h. The short target DNA (58 nt) is doubly labeled with a Cy3 label (green) at the non-target strand (NTS) and a Cy5 label (red) at the target strand (TS). **A** Cy3 (green sphere) and Cy5 (red sphere) labels are attached to the 5’-end of the DNA strands. **B** The cleaved NTS shows a different cleavage pattern for the various Cas12a variants. The TS cleavage pattern is the same for all Cas12a variants. **C** Cy3 (green sphere) and Cy5 (red sphere) labels positioned at the 3’-end of the DNA strands. **D** The cleavage patterns of target and non-target strand are the same for all Cas12a variants. Samples were analyzed on a high-resolution denaturing 15% PAA gel.

The same target DNA construct carrying the fluorescent labels at the 3’-ends was used to analyze the cleavage pattern on the PAM-proximal part. We found a comparable cleavage pattern for all Cas12a variants (Figure 2B). The 3’-labeled TS is processed yielding three products as already observed for the 5’-labeled DNA variant. Kinetic analysis shows that no trimming takes place at the PAM-distal part of the TS (Figure S5D/F). In agreement with data by Swarts *et al*., we detected two cleavage products for the NTS that simultaneously appear after a very short cleavage time (2). This suggests an initial imprecise cleavage at two different positions, before the trimming of the PAM-proximal part takes place (19). These two initial cleavage sites were detected for all Cas12a bridge helix mutants (Figure 2D). Taken together, mutations of the BH shift the final cleavage position of Cas12a on the NTS towards the 3’ end of the DNA strand and the BH is crucially involved in the trimming activity of Cas12a.

### Cas12a adopts three conformational states during its activity cycle

A number of X-ray and cryo-EM structures are available for Cas12a that show the architecture of binary and ternary Cas12a complexes(2, 17, 36–38, 20, 21, 30–35) and elucidated that a transition from the binary to ternary complex is accompanied by a structural re-arrangement from a closed to a more open state of the enzyme (Figure S7). However, no structural information is available for the apo Cas12a enzyme and for the pre-crRNA-Cas12a complex in the absence of a DNA target. In order to follow the conformational transitions of Cas12a and to evaluate whether the BH mutations affect the conformation of Cas12a, we performed smFRET measurements on diffusing molecules. To determine relative intramolecular distances in WT Cas12a, we site-specifically incorporated the unnatural amino acid *para*-azido-L-phenylalanine (AzF) via the Amber-suppressor strategy (39, 40). We placed the label either in the WED domain (K647AzF), the Nuc (T1222AzF) or REC lobe (D470AzF) yielding the following double mutants: Cas12a^K647AzF/T1222AzF^ (termed Cas12a^WED-Nuc^), Cas12a^D470AzF/K647AzF^ (referred to Cas12a^WED- REC^), and Cas12a^D470AzF/T1222AzF^ (referred to Cas12a^REC-Nuc^) (Figure S7). Coupling of the fluorescent dyes DyLight550 and DyLight650 via the azide group of AzF was performed via the Staudinger-Bertozzi ligation(41) (Figure S8). We used a stochastic labelling approach that yields proteins that carry a donor and acceptor dye pair or proteins that carry only the donor or acceptor. Cas12a molecules that carry the donor-acceptor pair can be spectroscopically be sorted based on the (PIE) excitation scheme(42) with separate detection channels for donor and acceptor emission. Crystal structures show the following distances between the chosen residues in the binary and ternary complex, respectively: positions K647 and T1222: 78.1 Å and 79.8 Å, D470 and K647: 98.0 Å and 104.4 Å, and D470 and T1222 30.8 Å and 36.0 Å, respectively (2, 21) (Figure S7). EMSAs and activity assays demonstrated that all mutants were able to bind nucleic acids and to cleave the target DNA (Figure S4, Figure S9).

First, we used the Cas12a^WED-Nuc^ and Cas12a^WED-REC^ variants to follow conformational changes of Cas12a along the WED-Nuc lobe axis and the WED-REC lobe axis, respectively. To this end, we performed smFRET measurements using (i) the apo enzyme, (ii) the binary complex composed of Cas12a plus pre-crRNA, (iii) the binary complex composed of Cas12a plus mature crRNA, (iv) the ternary complex (Cas12a + mature crRNA + dsDNA target) and (v) a ternary complex loaded with a single-stranded target. The Cas12a^WED-REC^ and Cas12a^WED-Nuc^ variants showed a single low FRET population for the apo enzyme, the binary and both variants of the ternary complex, respectively (Figure 3B). This indicates that the distance between WED and Nuc domain and the WED domain and REC lobe do not significantly change when Cas12a progresses through its activity cycle. This is in agreement with structural studies (2, 21) and molecular dynamics simulations (43), in which only marginal changes in distances between these positions can be detected (Figure S7). Using the Cas12a^Nuc-REC^ variant, a low FRET population with a FRET efficiency of 0.12 (± 0.04) and a high FRET population (E = 0.97 ± 0.01) can be detected for the nucleic-acid free Cas12a. This indicates that the apo form can adopt a closed and open conformation with the majority of the molecules (68 ± 7%) found in the closed state. Addition of crRNA to the protein shifts the molecules to the high FRET population (84 ± 2%). This effect is noticeable after the addition of mature crRNA, but also upon addition of pre-crRNA (Figure S10A). Cleavage assays using samples that contained pre-crRNA or the mature crRNA (Figure S10D/E) yielded comparable cleavage efficiencies. This demonstrates that the pre-crRNA is processed by Cas12a as only the mature crRNA allows loading and cleavage of dsDNA (16). Even though we cannot rule out that the pre-crRNA is partly processed during the incubation and single-molecule measurement time, the data nevertheless indicate that the pre-crRNA/Cas12a complex does not give rise to an additional FRET population and hence, it can be assumed that the pre-crRNA/Cas12a complex does adopt a distinct conformation. Notably, the formation of the binary complex is dependent on the crRNA concentration added to Cas12a underscoring the observation that the high FRET population reflects the binary complex (Figure S11A). crRNA-binding therefore induces the closure of the protein reflecting the closed state of Cas12a(2) (Figure S7). Upon loading of the DNA, a third population with a FRET efficiency of 0.82 (± 0.02) emerges. This indicates that the protein re-opens thereby providing enough space for the binding of target DNA. The relative number of molecules in the middle FRET population increased while the number of molecules that adopt the high FRET state decreased with increasing target DNA concentration indicating that the loading of DNA triggers the emergence of the middle FRET population (Figure S11B). The measurements were also performed with the catalytic inactive variant of Cas12a (dCas12a, Figure S12). Here, highly comparable data were obtained. This demonstrates that the medium FRET population corresponds to the ternary complex with bound target DNA and that the conformations of the ternary complex before cleavage (dCas12a) and after cleavage (WT Cas12a) of the target DNA do not significantly differ. Interestingly, loading is only moderately efficient and approximately 30-50% of the proteins loaded with a crRNA remain in the closed FRET state (Figure S11B). Additionally, we probed a ternary complex that lacks the NTS. Cas12a can process the TS even in the absence of the NTS (Figure 3, Figure S13) (20). In the absence of the NTS, the high FRET population drops to 9% (in the presence of NTS: 40%) and the middle FRET population increases to 60% indicating that loading of the sterically less demanding single-stranded TS is very efficient allowing a successful transition from the binary to the ternary complex. The FRET efficiency of the middle FRET population, however, does not significantly change suggesting that loading of the TS is sufficient to induce the opening of DNA cleft. Taken together, we show that the smFRET measurements re-capitulate the structural data of the binary complex (closed state) and the opening movement of Cas12a upon target DNA loading (semi-closed state). A single-stranded TS is sufficient and more efficient to induce the transition into the semi-closed state. Moreover, we can assess the structural state of the apo enzyme and show that apo Cas12a can adopt an open conformation.

**Figure 3.**
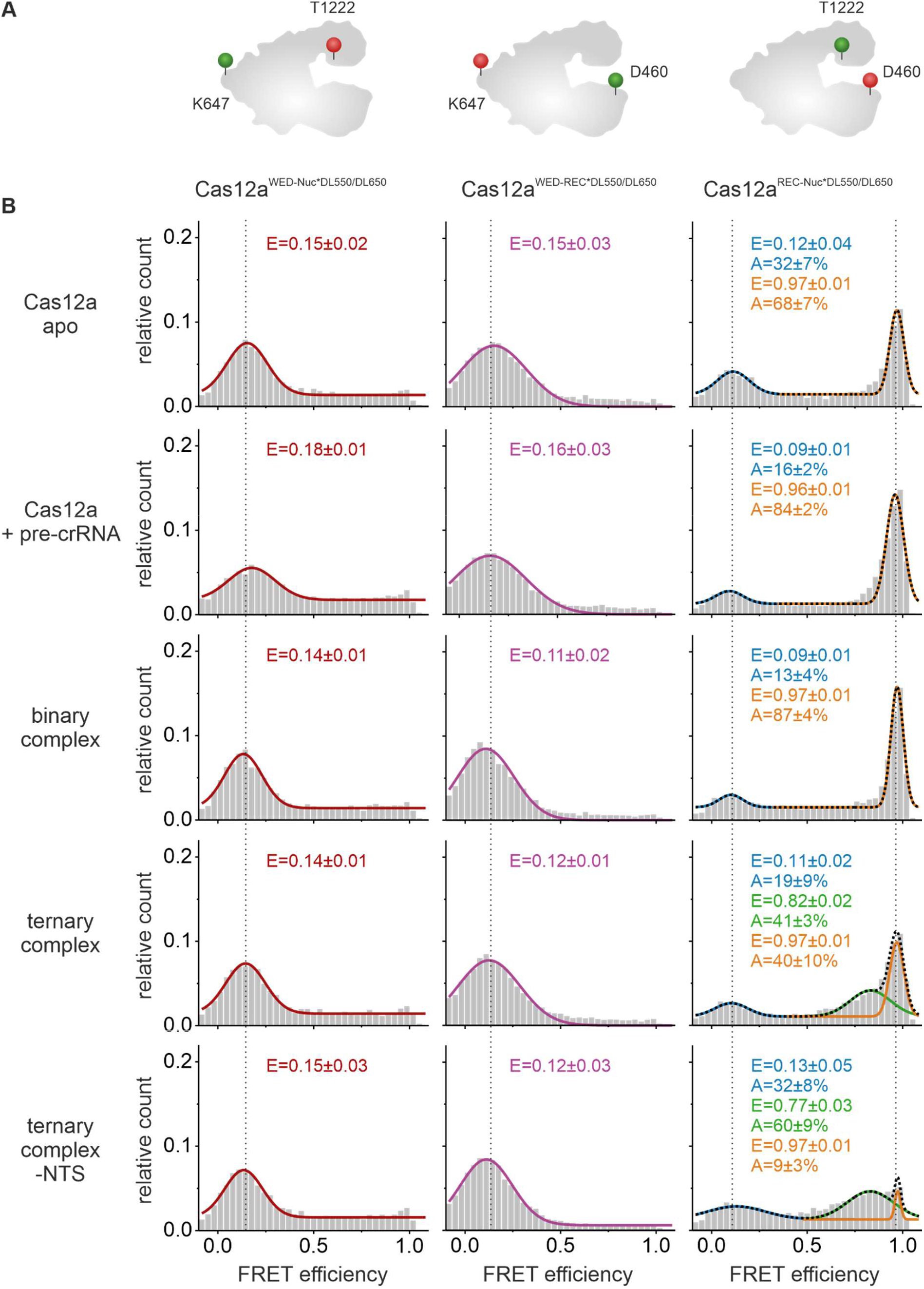
Fluorescence energy transfer (FRET) measurements reveal conformational states of Cas12a. Single-molecule FRET measurements using doubly labelled Cas12a variants were performed on freely diffusing molecules. **A** Schematic model of Cas12a showing the labeling positions at D470 (REC domain), K647 (WED domain), and T1222 (Nuc domain). **B** FRET efficiency distributions for the Cas12a apo enzyme, the Cas12a with pre-crRNA (1 nM), the binary complex (1 nM crRNA), the ternary complex (1 nM crRNA, 1 nM target DNA), and the ternary complex without NTS (1 nM crRNA, 1 nM TS DNA). Cas12a^WED-Nuc*DL550/DL650^ and Cas12a^WED-REC*DL550/DL650^ display one single FRET population for the apo, RNA-and RNA-and-DNA-bound state at a FRET efficiency of 0.15 ± 0.01 and 0.13 ± 0.02, respectively. Cas12a^REC-Nuc*DL550/DL650^ shows two FRET populations for the apo complex and the RNA-bound state at 0.11 ± 0.02 and 0.97 ± 0.01 with the high FRET population increasing with the addition of RNA. In the ternary complex, Cas12a^REC-Nuc*DL550/DL650^ shows three FRET populations with an additional population at 0.82 ± 0.02 FRET efficiency. Histograms show the data of three independent measurements. The histograms were fitted with a single, double or triple Gaussian function and the mean FRET efficiencies (E) and the percentage distribution of the populations (A) are given with SEs in the histograms.

### An intact BH is required for efficient binary complex formation

Being able to follow the opening and closure of the protein using the Cas12a^Nuc-REC^ variant, we additionally interrogated all BH mutants to understand whether the BH is involved in the stabilization of the conformational states of Cas12a (Figure 4). For all BH variants, low, middle and high FRET populations were detected. However, differences in the relative distribution of these states were observed. In the apo state, all W971 mutants showed a shift towards the low FRET population suggesting that a destabilization or modulation of the anchoring of the BH in the REC lobe increases the probability to adopt the open state. Interestingly, in case of the ΔBH mutant, the FRET efficiencies of the respective populations did not change and high and low FRET states were almost equally populated. As the residue W971 is still present in this mutant, we hypothesized that this might be sufficient to maintain the opened and closed state. In order to test this hypothesis, we made use of the so-called ΔΔBH in which not only residues 953-969 of the BH but in addition residues 970/971 were deleted resulting in a Cas12a variant fully devoid of the BH. Here, the relative distribution of the FRET states and the FRET efficiencies were not significantly altered (Figure S14). For all Cas12a variants, no dynamic switching between the open and closed state was observed during the millisecond timescale that we capture with our measurements. We conclude that the BH is not the primary structural element that regulates the open and closed state of Cas12a.

**Figure 4.**
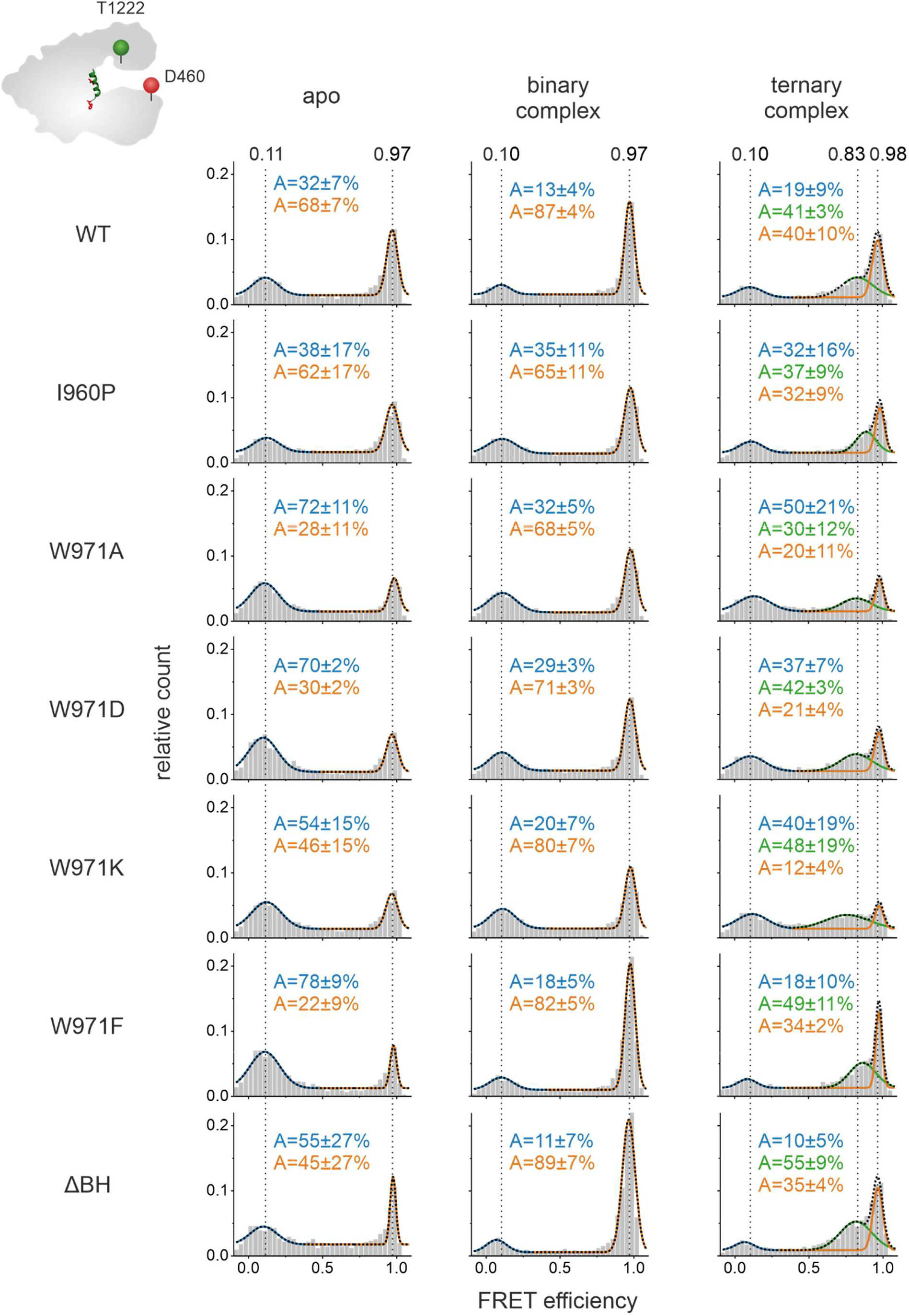
Fluorescence energy transfer (FRET) measurements reveal conformational states of Cas12a bridge helix mutants. Single-molecule FRET measurements on freely diffusing molecules were performed using the doubly labelled Cas12a^REC-Nuc*DL550/DL650^ variant that carried additional mutations in the bridge helix. FRET efficiency distributions for the apo enzyme, the binary complex (1 nM crRNA), and the ternary complex (1 nM crRNA, 1 nM target DNA) are shown. The apo histograms exhibits two FRET populations corresponding to an open and closed state of Cas12a, respectively. The addition of crRNA leads to an increase of the high FRET population in the binary complex. Addition of crRNA and target DNA give rise to a third population (semi-closed conformation). The histograms were fitted with a single or double Gaussian function and the averaged mean FRET efficiencies (E) and the percentage distribution of the populations (A) are given with SEs in the histograms. FRET efficiencies for the single protein variants are listed in **table S2**.

**Figure 5.**
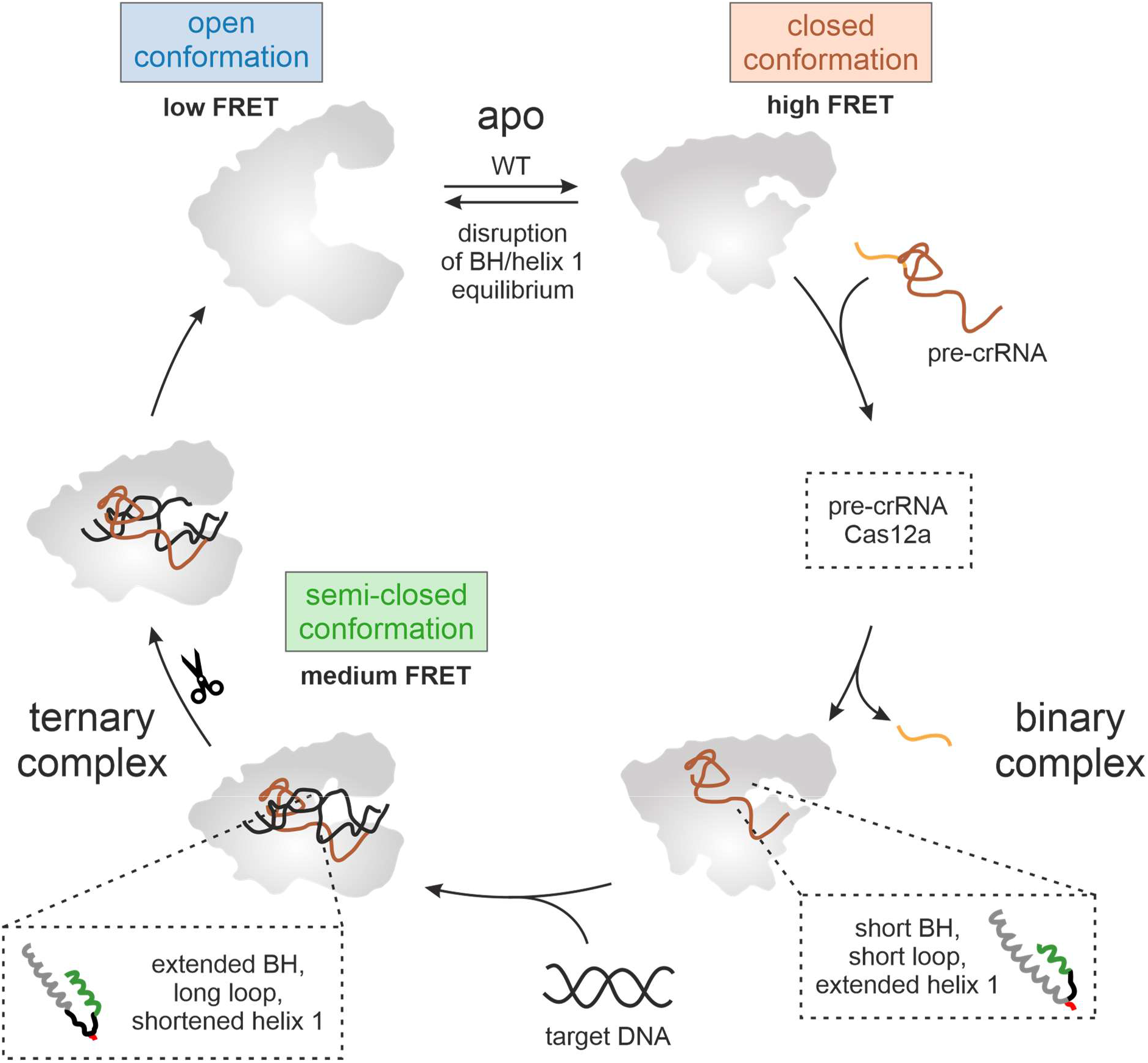
Impact of the bridge helix on the conformational flexibility and catalytic activity of Cas12a. In the apo state, FnCas12a can adopt two conformations: a closed (high FRET population) and open state (low FRET population). Binding of (pre-)crRNA induces the closure of the enzyme. In order to bind a single-stranded or double-stranded target DNA, Cas12a re-opens again giving rise to a medium FRET population in single-molecule FRET measurements. The bridge helix (BH, in green) is a central structural element in Cas12a that connects the REC and Nuc lobe with the tethering residue tryptophan 971 (shown in red) that anchors the BH in a hydrophobic pocket in the REC lobe. The length of the BH (green), the adjacent helix 1 (grey) and the interconnecting linker (black) as well as the position of W971 re-adjust upon transition from the binary to the ternary complex (see insets). Mutations in the BH or replacement of W971 affect trimming activity of Cas12a and influence the equilibrium between open and closed state of Cas12a.

All mutants adopt the closed conformation once the crRNA is loaded into the enzyme. crRNA titration experiments revealed, however, that the formation of the high FRET population is less efficient for mutants I960P, W971A/D/K (Figure S11A). In contrast, mutant W971F and the ΔBH mutant readily formed the closed state upon crRNA addition even at low crRNA concentrations (Figure S11A). If a mutant preferentially adopted the open state in the apo form, the loading of crRNA was inefficient. This suggests that for a subset of BH mutants binding of the crRNA or the conformational transition from the open to the closed state is impaired resulting in reduced amounts of the binary complex (Figure S15). In contrast, formation of the ternary complex is efficiently induced upon target DNA addition in all BH mutants (Figure 4 and S11B) implying that the mutations in the BH do not affect the loading of the target DNA. Please note that in all Cas12a variants, the middle FRET population shows relatively broad distribution suggesting that this state is more flexible as compared to the high FRET state (Figure S16). Taken together, the single-molecule FRET data indicate that a subset of BH mutants is not only impaired in their cleavage activity but that changes in the BH also disturb efficient binary complex formation.

## Discussion

The single effector nucleases Cas9 and Cas12a of the CRISPR-Cas type II system form a characteristic bilobal structure comprised of the REC and Nuc lobe, which are connected by the bridge helix. In Cas9, the BH reaches deep into the REC lobe and contacts directly or indirectly the target DNA, crRNA and tracrRNA (44–49). An arginine-rich patch in the BH orientates the nucleotides of the seed region of the crRNA to render them solvent-exposed. Thus, the crRNA can readily base-pair with the target DNA strand. Consequently, mutations in the BH of Cas9 of *Streptococcus pyogenes* (SpyCas9) influence RNA and target DNA binding, mismatch sensing and RNA-loop stability (49–52). Several SpyCas9 BH mutants showed increased specificity rendering them more suitable for genetic engineering purposes (51). The BH of FnCas12a also connects the REC and Nuc lobe but does not insert as deeply into the REC lobe as observed for SpyCas9 (2, 21). The BH of FnCas12a is not in close contact with the target DNA or crRNA and only contacts the sugar-phosphate backbone of the crRNA (31). The seed region of the crRNA is arranged by the REC1 domain instead. Nonetheless, our mutational analysis shows that the BH of Cas12a contributes to the catalytic activity and specificity of the enzyme. Deletion of the entire BH severely reduced the catalytic activity. Similarly, a BH deletion mutant of SpyCas9 was inactive (53). This shows that the BH is an essential segment for both enzymes. Additionally, we evaluated whether the helical nature of the BH or the anchoring of the BH in the REC domain is a pre-requisite for full catalytic activity. Introduction of a proline residue at position 960 in the BH resulted in a reduced NTS-trimming activity showing that the helical nature is required to reach the full activity spectrum of Cas12a. Disturbing the anchoring of the BH in the REC domain affected the cleavage reaction rate and/or trimming activity. Charged amino acids led to a reduction of the DNA cleavage rate. Additionally, the trimming activity of Cas12a was also lost. In contrast, an alanine substitution did not affect the cleavage and trimming activity. Replacement of W971 by a phenylalanine residue only marginally influenced the cleavage activity. Trimming of the NTS was only slightly affected. The differing effect of the mutations can be rationalized from a biochemical perspective: charged residues cause a repulsion of the BH from the hydrophobic pocket in the REC2 domain whereas the small and uncharged alanine substitution might still fit in the pocket. Phenylalanine is also a hydrophobic residue and can most likely functionally replace tryptophan. Consequently, the mutant is only mildly affected. Similar to some mutants evaluated in this work, Swarts *et al*. identified Cas12a mutants that exhibited reduced trimming activity (e.g. R918, D920, S1083, R1218, D1255, D1227). However, these mutants are constituting the active site or are in close contact to the catalytic residues. This is not the case for the mutants tested in this study. In order to rationalize the observed trimming defects based on structural data, we carefully inspected all Cas12a structures available. Strikingly, we found that the BH length expands upon ternary complex formation. Notably, structural changes in the BH occur in tandem with structural changes in the adjacent helix 1 of the RuvC domain (Figure S17). The re-positioning of the binding pocket in the REC domain requires the positional shift of tryptophan 971 by 8.7 Å to allow docking of the residue in its binding pocket also in the ternary complex (Figure S17A/B). To promote this movement of W971, the residue has to be released from helix 1 as observed in ternary complex structures that show that helix 1 is shortened by six residues thereby releasing W971 into a flexible loop that connects the BH and helix 1. Concomitantly, the BH is extended by approximately 2 residues and 4 residues in FnCas12a and LbCas12a, respectively. Hence, these two helices act in concert as “suspension helices” to compensate for the large conformational changes of the REC domains upon transition to the ternary complex. Part of the BH is not resolved in the binary complex structure indicating that this is a flexible region in the Cas12a-crRNA complex. Furthermore, a conserved lysine residue (K978 in FnCas12a) in helix 1 contacts the backbone of the target DNA in the ternary complex. Due to a shift in the position of helix 1 upon transition to the ternary complex, K978 is ideally orientated for the interaction with the DNA. While K978 is positioned in the middle of helix 1 in the binary complex, it is shifted to the N-terminal end of the re-modelled helix 1 and not released into the flexible loop in the ternary complex. This indicates that the integration of K978 into helix 1 is essential to function as DNA-contacting base. It seems likely that some of the mutations introduced into the BH of FnCas12a affect the tandem movement of the BH and helix 1 and consequently, DNA contacts and the position of the REC domains are slightly altered. Altogether, these disruptions most likely cause the observed trimming defect. For example, formation of an extended BH in the ternary complex might be prevented by the I960P mutant that is located in the middle of the BH. Similarly, the absence of the entire BH completely disrupts the coupled helical restructuring events. In addition to repulsion of the W971D or W971K residues from the hydrophobic binding pocket, these substitutions might also affect the formation of the extended BH as position 971 re-positions to the very beginning of the loop (next to the end of the BH). Notably, the tandem helix arrangement is conserved in all Cas12a variants studied so far (Figure S17).

Crystal and cryo-EM structures (2,17,36–38,20,21,30–35) as well as smFRET studies (20, 23, 54) indicated that not only the protein itself but also nucleic acids undergo substantial conformational changes. Hence, we employed smFRET measurements to follow the movement of the REC and Nuc lobe of Cas12a WT and the BH mutant variants. Our data faithfully replicate the structural states of the binary and ternary complex known from crystal structures and add hitherto missing information about the conformational state of the apo enzyme. More specifically, we observe a closure of the Cas12a protein upon loading of the crRNA and a slight re-opening upon loading of the DNA. This is in agreement with structural data for FnCas12a and AsCas12a (2,17,36–38, 20,21,30–35). Our finding that the ternary complex is more flexible than the binary complex agrees with recent molecular dynamics simulations that showed the increased flexibility of the REC and Nuc lobe in the ternary complex (43). We found that loading of the DNA target is more efficient when a ssDNA target is provided suggesting that the formation of a Cas12a-crRNA-dsDNA complex is energetically demanding. In line with crystal structures, the expansion of REC and Nuc lobe upon transition to the ternary complex does not depend on the single-stranded or double-stranded nature of the DNA and loading of the ts is sufficient to induce the conformational change in Cas12a (21). Conformational heterogeneity was detected for the apo enzyme. Here, Cas12a can adopt an open or closed state. No crystal or high resolution cryo-EM structure of the apo enzyme is available so far. But the existence of an open state has been proposed based on smFRET studies by Stella *et al*. While we performed smFRET measurements on diffusing molecules, Stella *et al*. employed immobilized Cas12a proteins to follow the relative distance between the REC and Nuc lobe (20). The apo enzyme and all complexes tested in the study by Stella *et al*. showed a wide range of conformations that could not be discerned as discrete states. However, in the apo state, a relatively high fraction of the molecules was found in a low FRET state supporting the finding of an open state in this study. Considering the central position of the BH, it seemed feasible that a deletion of the BH might result in a collapsed state of Cas12a. However, this was not observed in our single-molecule FRET measurements. Interestingly, the structural integrity is preserved even in the ΔBH/ΔΔBH variants and these mutants can load crRNA and target DNA efficiently. Considering the previous finding that the BH structurally re-arranges in concert with helix 1 of the RuvC I domain it seems plausible that an intact helix 1 (possibly in addition to structural elements in the PI and Wedge domains) warrants the correct distance between the REC and Nuc lobe. Consequently, the BH only influences the equilibrium between the open and closed state in the apo enzyme. Additional structural data that elucidate the structural state of the BH and helix 1 in the apo enzyme would be of high interest in this context.

Noteworthy, the mutants that cause a reduced trimming activity produce a more precise cleavage pattern as cleavage occurs mainly at two sites. In contrast, the WT enzyme produces at least five intermediate products on the way to the two final products. While crRNA loading seems to occur with reduced efficiency in these mutants, they do not affect the conformational changes required for the catalytic cycle of Cas12a. Hence, these mutants might be better suited for gene editing applications than the WT as they might warrant a more precise cleavage pattern.

## Data availability

The datasets generated during and/or analyzed during the current study are available from the corresponding author on reasonable request.

## Acknowledgements

We gratefully thank Dr. Sarah Willkomm and Kevin Kramm for fruitful discussions and Elisabeth Piechatschek and Elke Papst for technical assistance.

## Author contributions

D.G. conceived the study. E.W., L.J. and A.S. performed the single-molecule measurements. E.W., L.J. and A.S. analyzed the single-molecule data. E.W., L.J., A.S. and G.Z. purified the proteins and created Cas12a mutants. E.W., L.J., A.S. and G.Z. performed EMSAs and cleavage assays. E.W. and D.G. wrote the paper. All authors commented on the paper.

## References

1. Zetsche, B., Gootenberg, J.S., Abudayyeh, O.O., Slaymaker, I.M., Makarova, K.S., Essletzbichler, P., Volz, S.E., Joung, J., Van Der Oost, J., Regev, A., et al. (2015) Cpf1 Is a Single RNA-Guided Endonuclease of a Class 2 CRISPR-Cas System. Cell, 163, 759–771.

2. Swarts, D.C., van der Oost, J. and Jinek, M. (2017) Structural Basis for Guide RNA Processing and Seed-Dependent DNA Targeting by CRISPR-Cas12a. Mol. Cell, 66, 221-233.e4.

3. Makarova, K.S., Wolf, Y.I., Iranzo, J., Shmakov, S.A., Alkhnbashi, O.S., Brouns, S.J.J., Charpentier, E., Cheng, D., Haft, D.H., Horvath, P., et al. (2019) Evolutionary classification of CRISPR–Cas systems: a burst of class 2 and derived variants. Nat. Rev. Microbiol., 18.

4. Nuñez, J.K., Kranzusch, P.J., Noeske, J. and Wright, A. V (2014) Cas1 – Cas2 complex formation mediates spacer acquisition during CRISPR – Cas adaptive immunity. Nat. Struct. Mol. Biol., 21, 528–534.

5. Amitai, G. and Sorek, R. (2016) CRISPR-Cas adaptation: Insights into the mechanism of action. Nat. Rev. Microbiol., 14, 67–76.

6. Mohanraju, P., Makarova, K.S., Zetsche, B., Zhang, F., Koonin, E. V. and van der Oos, J. (2016) Diverse evolutionary roots and mechanistic variations of the CRISPR-Cas systems. Science, 353.

7. Charpentier, E., Richter, H., van der Oost, J. and White, M.F. (2015) Biogenesis pathways of RNA guides in archaeal and bacterial CRISPR-Cas adaptive immunity. FEMS Microbiol. Rev., 39, 428–441.

8. Hille, F., Richter, H., Wong, S.P., Bratovič, M., Ressel, S. and Charpentier, E. (2018) The Biology of CRISPR-Cas: Backward and Forward. Cell, 172, 1239–1259.

9. Jinek, M., Chylinski, K., Fonfara, I., Hauer, M., Doudna, J.A. and Charpentier, E. (2012) A programmable dual-RNA-guided DNA endonuclease in adaptive bacterial immunity. Science (80-.)., 337, 816–821.

10. Zetsche, B., Heidenreich, M., Mohanraju, P., Fedorova, I., Kneppers, J., Degennaro, E.M., Winblad, N., Choudhury, S.R., Abudayyeh, O.O., Gootenberg, J.S., et al. (2017) Multiplex gene editing by CRISPR-Cpf1 using a single crRNA array. Nat. Biotechnol., 35, 31–34.

11. Knott, G.J. and Doudna, J.A. (2018) CRISPR-Cas guides the future of genetic engineering. Science (80-.)., 361, 866–869.

12. Barrangou, R. and Doudna, J.A. (2016) Applications of CRISPR technologies in research and beyond. Nat. Biotechnol., 34, 933–941.

13. Pickar-Oliver, A. and Gersbach, C.A. (2019) The next generation of CRISPR–Cas technologies and applications. Nat. Rev. Mol. Cell Biol., 20, 490–507.

14. Shmakov, S., Smargon, A., Scott, D., Cox, D., Pyzocha, N., Yan, W., Abudayyeh, O.O., Gootenberg, J.S., Makarova, K.S., Wolf, Y.I., et al. (2017) Diversity and evolution of class 2 CRISPR–Cas systems. Nat. Rev. Microbiol., 15, 169–182.

15. Nishimasu, H. and Nureki, O. (2017) Structures and mechanisms of CRISPR RNA-guided effector nucleases. Curr. Opin. Struct. Biol., 43, 68–78.

16. Fonfara, I., Richter, H., Bratovič, M., Le Rhun, A. and Charpentier, E. (2016) The CRISPR-associated DNA-cleaving enzyme Cpf1 also processes precursor CRISPR RNA. Nature, 532, 517–521.

17. Yamano, T., Zetsche, B., Ishitani, R., Zhang, F., Nishimasu, H. and Nureki, O. (2017) Structural Basis for the Canonical and Non-canonical PAM Recognition by CRISPR-Cpf1. Mol. Cell, 67, 633-645.e3.

18. Specht, D.A., Xu, Y. and Lambert, G. (2020) Massively parallel CRISPRi assays reveal concealed thermodynamic determinants of dCas12a binding. Proc. Natl. Acad. Sci., 10.1073/PNAS.1918685117.

19. Strohkendl, I., Saifuddin, F.A., Rybarski, J.R., Finkelstein, I.J. and Russell, R. (2018) Kinetic Basis for DNA Target Specificity of CRISPR-Cas12a. Mol. Cell, 71, 816-824.e3.

20. Stella, S., Mesa, P., Thomsen, J., Paul, B., Alcón, P., Jensen, S.B., Saligram, B., Moses, M.E., Hatzakis, N.S. and Montoya, G. (2018) Conformational Activation Promotes CRISPR-Cas12a Catalysis and Resetting of the Endonuclease Activity. Cell, 175, 1856-1871.e21.

21. Swarts, D. and Jinek, M. (2018) Mechanistic Insights into the Cis-and Trans-acting Deoxyribonuclease Activities of Cas12a. Mol. Cell, 73, 1–12.

22. Cofsky, J.C., Karandur, D., Huang, C.J., Witte, I.P., Kuriyan, J. and Doudna, J.A. (2020) CRISPR-Cas12a exploits R-loop asymmetry to form double strand breaks. Elife, 9, 2020.02.10.937540.

23. Jeon, Y., Choi, Y.H., Jang, Y., Yu, J., Goo, J., Lee, G., Jeong, Y.K., Lee, S.H., Kim, I.S., Kim, J.S., et al. (2018) Direct observation of DNA target searching and cleavage by CRISPR-Cas12a. Nat. Commun., 9.

24. Singh, D., Mallon, J., Poddar, A., Wang, Y., Tippana, R., Yang, O., Bailey, S. and Ha, T. (2018) Real-time observation of DNA target interrogation and product release by the RNA-guided endonuclease CRISPR Cpf1 (Cas12a). Proc. Natl. Acad. Sci., 115, 5444–5449.

25. Chen, J.S., Ma, E., Harrington, L.B., Da Costa, M., Tian, X., Palefsky, J.M. and Doudna, J.A. (2018) CRISPR-Cas12a target binding unleashes indiscriminate single-stranded DNase activity. Science (80-.)., 360, 436–439.

26. Li, S.Y., Cheng, Q.X., Li, X.Y., Zhang, Z.L., Gao, S., Cao, R.B., Zhao, G.P., Wang, J. and Wang, J.M. (2018) CRISPR-Cas12a-assisted nucleic acid detection. Cell Discov., 4, 18–21.

27. Gootenberg, J.S., Abudayyeh, O.O., Kellner, M.J., Joung, J., Collins, J.J. and Zhang, F. (2018) Multiplexed and portable nucleic acid detection platform with Cas13, Cas12a, and Csm6. Science (80-.)., 6387, 439–444.

28. Broughton, J.P., Deng, X., Yu, G., Fasching, C.L., Servellita, V., Singh, J., Miao, X., Streithorst, J.A., Granados, A., Sotomayor-Gonzalez, A., et al. (2020) CRISPR–Cas12-based detection of SARS-CoV-2. Nat. Biotechnol., 10.1038/s41587-020-0513-4.

29. Curti, L., Pereyra-Bonnet, F. and Gimenez, C.A. (2020) An ultrasensitive, rapid, and portable coronavirus SARS-CoV-2 sequence detection method based on CRISPR-Cas12. bioRxiv, 10.1101/2020.02.29.971127.

30. Dong, D., Ren, K., Qiu, X., Zheng, J., Guo, M., Guan, X., Liu, H., Li, N., Zhang, B., Yang, D., et al. (2016) The crystal structure of Cpf1 in complex with CRISPR RNA. Nature, 532, 522–526.

31. Yamano, T., Nishimasu, H., Zetsche, B., Hirano, H., Slaymaker, I.M., Li, Y., Fedorova, I., Nakane, T., Makarova, K.S., Koonin, E. V., et al. (2016) Crystal Structure of Cpf1 in Complex with Guide RNA and Target DNA. Cell, 165, 949–962.

32. Stella, S., Alcón, P. and Montoya, G. (2017) Structure of the Cpf1 endonuclease R-loop complex after target DNA cleavage. Nature, 546, 559–563.

33. Knott, G.J., Cress, B.F., Liu, J., Thornton, B.W., Lew, R.J., Al-Shayeb, B., Rosenberg, D.J., Hammel, M., Adler, B.A., Lobba, M.J., et al. (2019) Structural basis for AcrVA4 inhibition of specific CRISPR-Cas12a. Elife, 8, 1–25.

34. Dong, L., Guan, X., Li, N., Zhang, F., Zhu, Y., Ren, K., Yu, L., Zhou, F., Han, Z., Gao, N., et al. (2019) An anti-CRISPR protein disables type V Cas12a by acetylation. Nat. Struct. Mol. Biol., 26, 308–314.

35. Gao, P., Yang, H., Rajashankar, K.R., Huang, Z. and Patel, D.J. (2016) Type v CRISPR-Cas Cpf1 endonuclease employs a unique mechanism for crRNA-mediated target DNA recognition. Cell Res., 26, 901–913.

36. Peng, R., Li, Z., Xu, Y., He, S., Peng, Q., Wu, L., Wu, Y., Qi, J., Wang, P., Shi, Y., et al. (2019) Structural insight into multistage inhibition of CRISPR-Cas12a by AcrVA4. Proc. Natl. Acad. Sci., 116, 18928–18936.

37. Nishimasu, H., Yamano, T., Gao, L., Zhang, F., Ishitani, R. and Nureki, O. (2017) Structural Basis for the Altered PAM Recognition by Engineered CRISPR-Cpf1. Mol. Cell, 67, 139–147.e2.

38. Zhang, H., Li, Z., Daczkowski, C.M., Gabel, C., Mesecar, A.D. and Chang, L. (2019) Structural Basis for the Inhibition of CRISPR-Cas12a by Anti-CRISPR Proteins. Cell Host Microbe, 25, 815–826.e4.

39. Chin, J.W., Santoro, S.W., Martin, A.B., King, D.S., Wang, L. and Schultz, P.G. (2002) Addition of p-azido-L-phenylalanine to the genetic code of Escherichia coli. J. Am. Chem. Soc., 124, 9026–9027.

40. Grohmann, D., Werner, F. and Tinnefeld, P. (2013) Making connections-strategies for single molecule fluorescence biophysics. Curr. Opin. Chem. Biol., 17, 691–698.

41. Saxon, E. and Bertozzi, C.R. (2000) Cell surface engineering by a modified Staudinger reaction. Science (80-.)., 287, 2007–2010.

42. Hendrix, J. and Lamb, D.C. (2013) Pulsed interleaved excitation: Principles and applications 1st ed. Elsevier Inc.

43. Saha, A., Arantes, P.R., Narkhede, Y.B., Hsu, R. V., Jinek, M. and Palermo, G. (2020) DNA-Induced Dynamic Switch Triggers Activation of CRISPR-Cas12a. Submitted, 10.1021/acs.jcim.0c00929.

44. Yamada, M., Watanabe, Y., Gootenberg, J.S., Hirano, H., Ran, F.A., Nakane, T., Ishitani, R., Zhang, F., Nishimasu, H. and Nureki, O. (2017) Crystal Structure of the Minimal Cas9 from Campylobacter jejuni Reveals the Molecular Diversity in the CRISPR-Cas9 Systems. Mol. Cell, 65, 1109–1121.e3.

45. Jiang, F., Taylor, D.W., Chen, J.S., Kornfeld, J.E., Zhou, K., Thompson, A.J., Nogales, E. and Doudna, J.A. (2016) Structures of a CRISPR-Cas9 R-loop complex primed for DNA cleavage. Science (80-.)., 351, 867–871.

46. Jinek, M., Jiang, F., Taylor, D.W., Sternberg, S.H., Kaya, E., Ma, E., Anders, C., Hauer, M., Zhou, K., Lin, S., et al. (2014) Structures of Cas9 endonucleases reveal RNA-mediated conformational activation. Science (80-.)., 343.

47. Anders, C., Niewoehner, O., Duerst, A. and Jinek, M. (2014) Structural basis of PAM-dependent target DNA recognition by the Cas9 endonuclease. Nature, 513, 569–573.

48. Nishimasu, H., Cong, L., Yan, W.X., Ran, F.A., Zetsche, B., Li, Y., Kurabayashi, A., Ishitani, R., Zhang, F. and Nureki, O. (2015) Crystal Structure of Staphylococcus aureus Cas9. Cell, 162, 1113–1126.

49. Nishimasu, H., Ran, F.A., Hsu, P.D., Konermann, S., Shehata, S.I., Dohmae, N., Ishitani, R., Zhang, F. and Nureki, O. (2014) Crystal structure of Cas9 in complex with guide RNA and target DNA. Cell, 156, 935–949.

50. Zeng, Y., Cui, Y., Zhang, Y., Zhang, Y., Liang, M., Chen, H., Lan, J., Song, G. and Lou, J. (2018) The initiation, propagation and dynamics of CRISPR-SpyCas9 R-loop complex. Nucleic Acids Res., 46, 350–361.

51. Bratovič, M., Fonfara, I., Chylinski, K., Gálvez, E.J.C., Sullivan, T.J., Boerno, S., Timmermann, B., Boettcher, M. and Charpentier, E. (2020) Bridge helix arginines play a critical role in Cas9 sensitivity to mismatches. Nat. Chem. Biol., 16, 587–595.

52. Babu, K., Amrani, N., Jiang, W., Yogesha, S.D., Nguyen, R., Qin, P.Z. and Rajan, R. (2019) Bridge Helix of Cas9 Modulates Target DNA Cleavage and Mismatch Tolerance. Biochemistry, 10.1021/acs.biochem.8b01241.

53. Shams, A., Higgins, S.A., Fellmann, C., Laughlin, T.J., Lew, R., Lukarska, M., Arnold, M., Staahl, B.T. and Savage, D.F. (2020) Comprehensive deletion landscape of CRISPR-Cas9 identifies minimal RNA-guided DNA-binding modules 2 3. bioRxiv.

54. Zhang, L., Sun, R., Yang, M., Peng, S., Cheng, Y. and Chen, C. (2019) Conformational Dynamics and Cleavage Sites of Cas12a Are Modulated by Complementarity between crRNA and DNA. iScience, 19, 492–503.

55. Chin, J.W., Santoro, S.W., Martin, A.B., King, D.S., Wang, L. and Schultz, P.G. (2002) Addition of p-azido-L-phenylalanine to the genetic code of Escherichia coli. J. Am. Chem. Soc., 124, 9026–9027.

56. Creutzburg, S.C.A., Wu, W.Y., Mohanraju, P., Swartjes, T., Alkan, F. and Gorodkin, J. (2020) Good guide, bad guide?: Spacer sequence-dependent cleavage efficiency of. 10.1093/nar/gkz1240.

57. Schrimpf, W., Barth, A., Hendrix, J. and Lamb, D.C. (2018) PAM: A Framework for Integrated Analysis of Imaging, Single-Molecule, and Ensemble Fluorescence Data. Biophys. J., 114, 1518–1528.

58. Nir, E., Michalet, X., Hamadani, K.M., Laurence, T.A., Neuhauser, D., Kovchegov, Y. and Weiss, S. (2006) Shot-Noise Limited Single-Molecule FRET Histograms: Comparison between Theory and Experiments. J. Phys. Chem. B, 110, 22103–22124.

59. Hellenkamp, B., Schmid, S., Doroshenko, O., Opanasyuk, O., Kühnemuth, R., Rezaei Adariani, S., Ambrose, B., Aznauryan, M., Barth, A., Birkedal, V., et al. (2018) Precision and accuracy of single-molecule FRET measurements—a multi-laboratory benchmark study. Nat. Methods, 15, 669–676.

