## Supplementary Information for "Decoupling the bridge helix of Cas12a results in a reduced trimming activity and impaired conformational transitions"

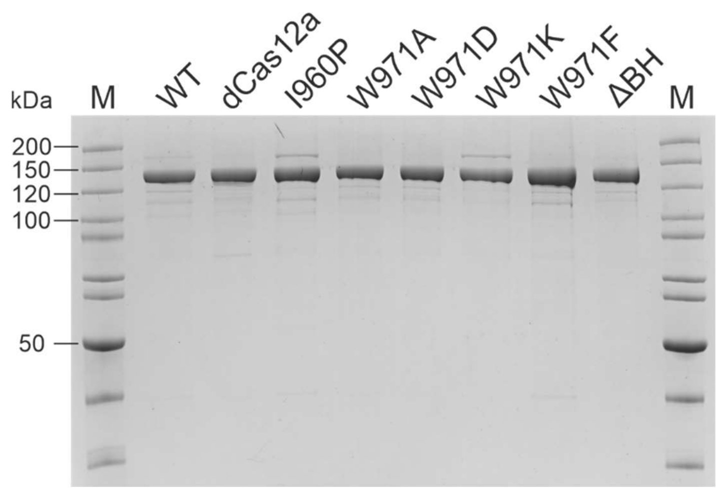


**Figure S1.** SDS-PAGE of purified FnCas12a variants (0.82 µg each, molecular weight of FnCas12a: 151.9 kDa). M: PageRuler™ (Thermo Scientific) unstained.

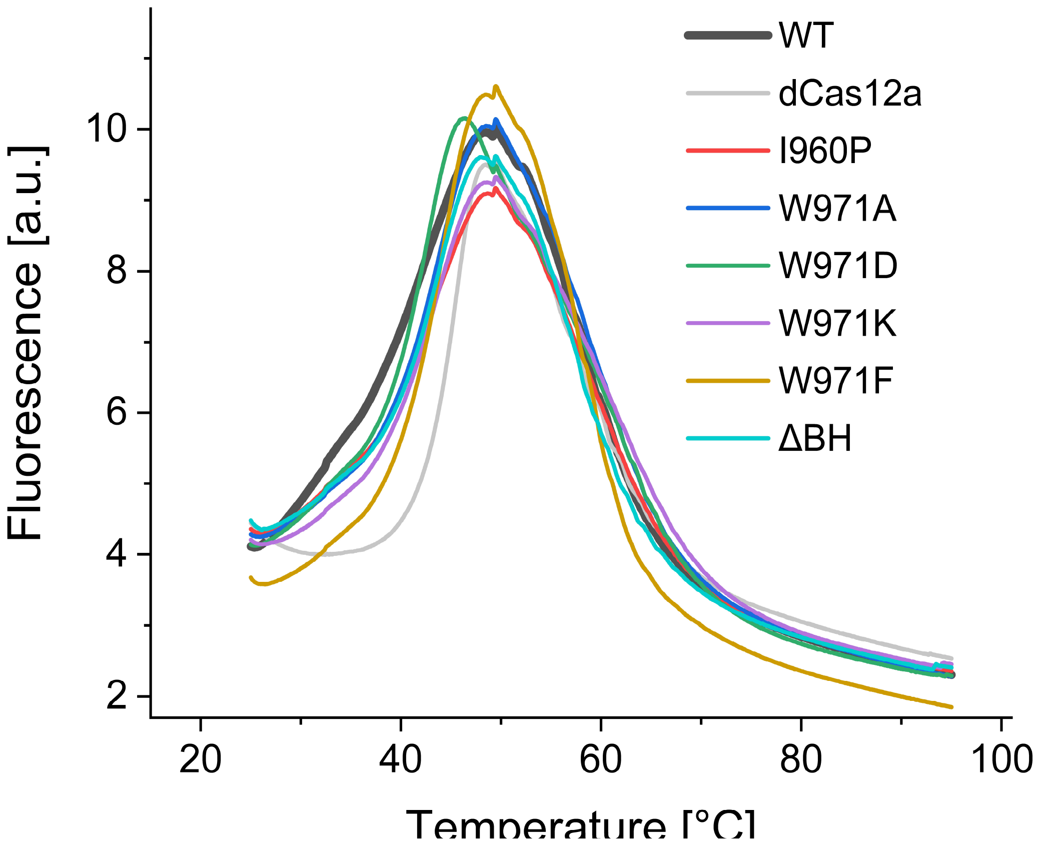


**Figure S2.** Melting curves of FnCas12a variants. Protein Thermal Shift™ (Thermo Scientific) melting curves of Cas12a variants (2 µg) from 25 to 95 °C, with an average $T_m = 48.2 \pm 0.7$ °C. Shown is the average of four replicates.

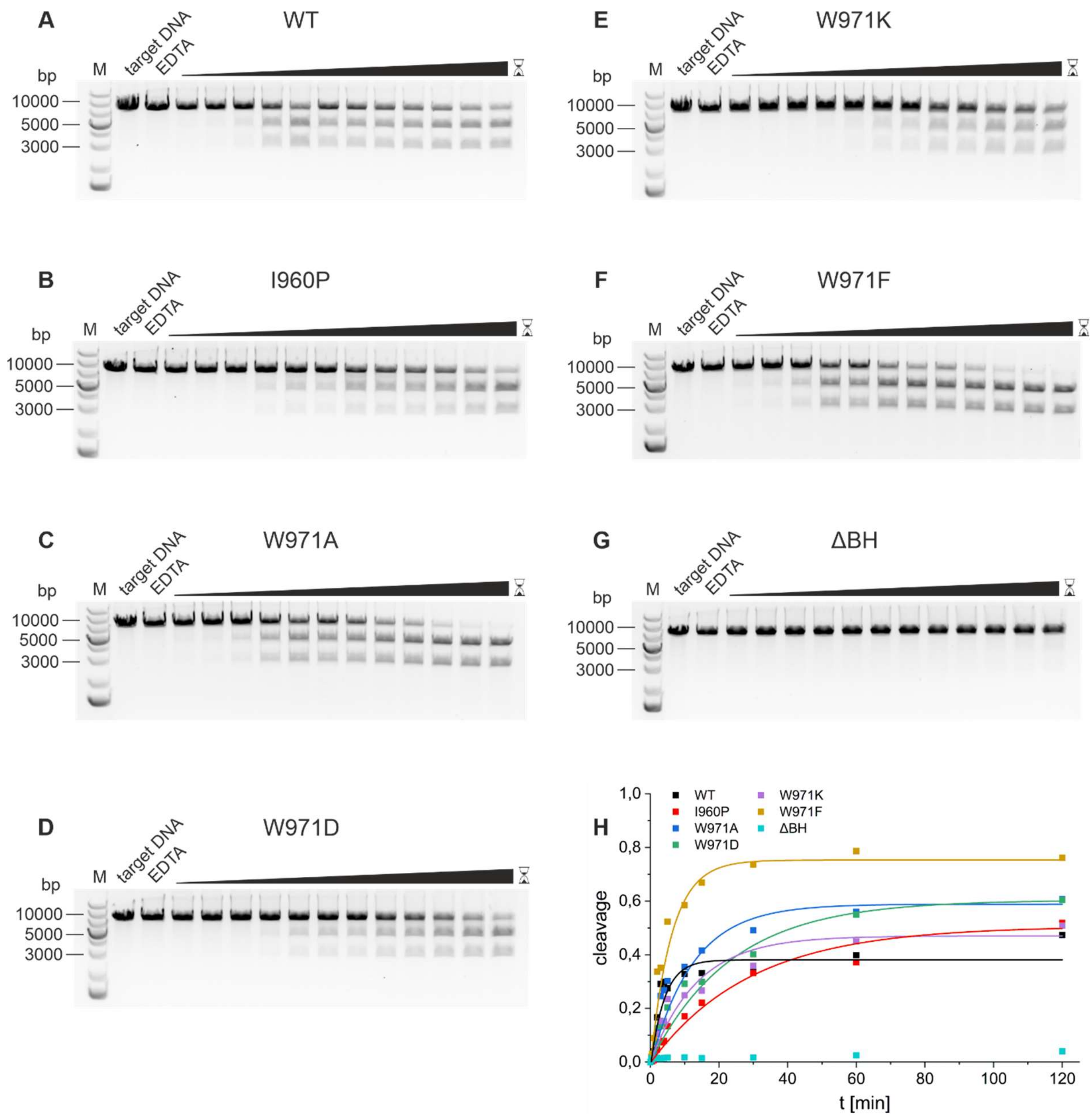


**Figure S3. Plasmid cleavage kinetics of WT Cas12a and Cas12a bridge helix mutants at 28 °C.** The reaction was composed of 25 nM Cas12a, 25 nM crRNA and 5 nM target DNA. Reactions were stopped by the addition of 83.3 mM EDTA after different time intervals of 15 s, 30 s, 1 min, 2 min, 3 min, 4 min, 5 min, 10 min, 15 min, 30 min, 1 h, and 2 h. M: 1 kb plus DNA ladder; target DNA: linearized target DNA without protein and crRNA; EDTA: reaction immediately stopped with EDTA after mixing of the reaction partners.

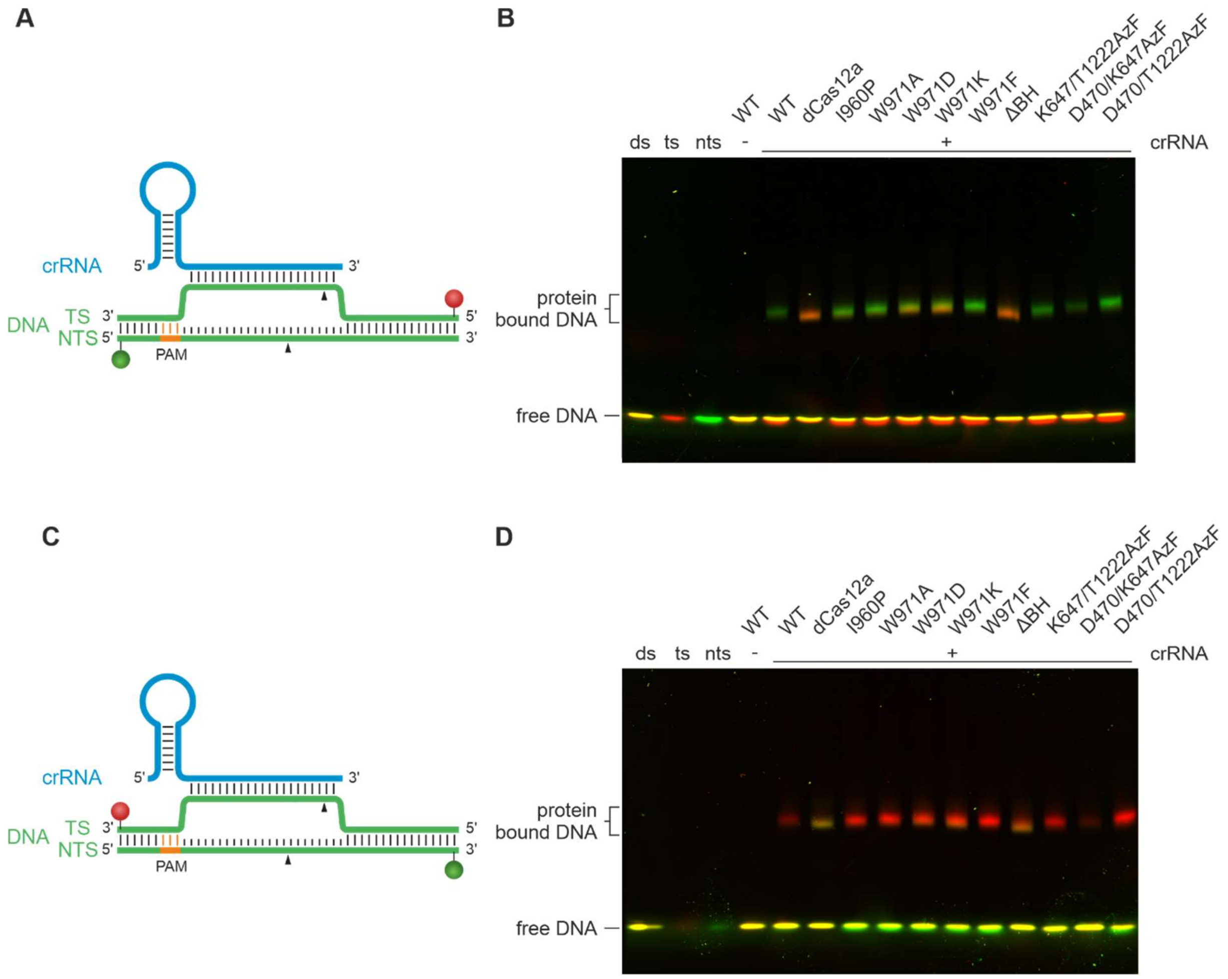


**Figure S4. Electrophoretic mobility shift assays of Cas12a-crRNA-target DNA complexes.** WT Cas12a, dCas12a, Cas12a bridge helix mutants, and Cas12a with incorporated AzF was used in a 7.5-fold excess of protein and crRNA (75 nM) over target DNA (10 nM). The short target DNA (58 nt) is doubly labeled with a Cy3 label (green) at the non-target strand (nts) and a Cy5 label (red) at the target strand (ts). **A** Cy3 (green sphere) and Cy5 (red sphere) labels are coupled to the 5'-end of the DNA strands, respectively. **B** Free DNA is detected at the bottom of the gel, protein-bound DNA is shifted and detected as slowly migrating complex. The protein-bound DNA band is mainly green, as the protein remains bound to the PAM proximal part of the DNA (Cy3) after DNA cleavage. dCas12a does not cleave the DNA and Cas12a$^{\Delta BH}$ is only slightly active, consequently, the Cy5-labeled DNA is still bound to the protein. **C** Cy3 (green sphere) and Cy5 (red sphere) labels are attached to the 3'-end of the DNA strands, respectively. **D** Free DNA is detected at the bottom of the gel, protein-bound DNA is detected as slowly migrating band. The protein-bound DNA band is mainly red because the protein remains bound to the PAM proximal part of the DNA (Cy5) after DNA cleavage. dCas12a does not cleave the DNA and Cas12a$^{\Delta BH}$ is only slightly active and consequently, the Cy3-labeled DNA is still bound to the protein. Please note that due to low protein expression yiedl, the Cas12a$^{D470/K647Azf}$ variant could only be used at a reduced concentration (60 nM) resulting in a reduced reduced fluorescence signal for the DNA-protein complex.

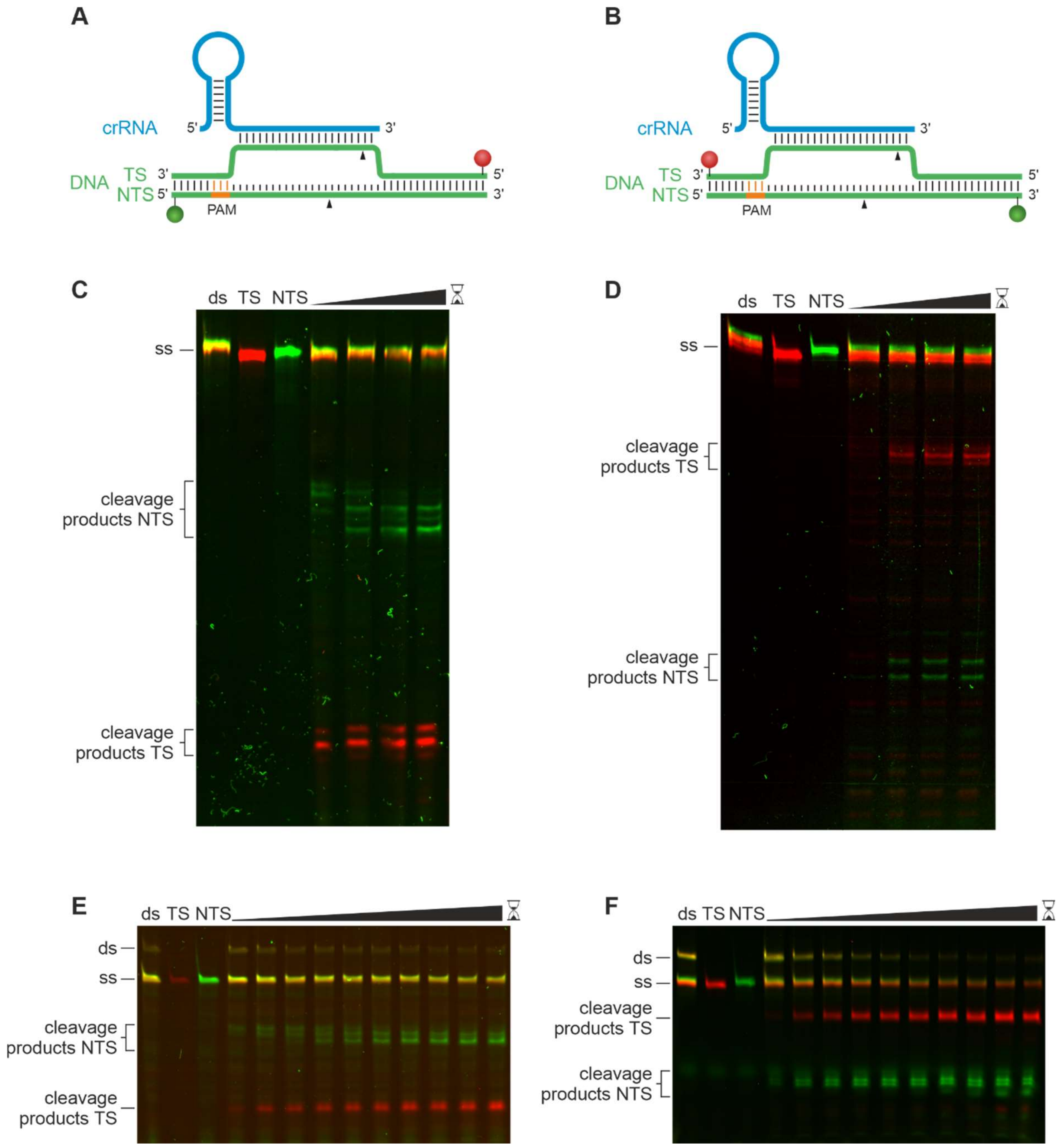


**Figure S5. Cleavage kinetics of WT Cas12a using a short doubly labeled DNA target.** Reactions contained Cas12a and crRNA (**C**, **D** 750 nM; **E**, **F** 75 nM) in 7.5-fold excess over the target DNA (58 nt; **C**, **D** 100 nM; **E**, **F** 10 nM). Reactions were stopped with 125 mM EDTA after different time intervals. **A** and **B** Schematic representation of the labeled DNA construct with crRNA. **A** Cy3 (green sphere) and Cy5 (red sphere) labels are coupled to the 5'-end of the DNA strands. **B** Cy3 (green sphere) and Cy5 (red sphere) labels are attached to the 3'-end of the DNA strands. **C** and **D** Cleavage kinetics on a high-resolution gel stopped after 1 s, 5 min, 30 min, and 1 h. **C** 5'-labeled target DNA; **D** 3'-labeled target DNA. **E** and **F** Cleavage kinetics stopped after 1 s, 30 s, 1 min, 3 min, 5 min, 10 min, 20 min, 30 min, 45 min, and 1 h. **E** 5'-labeled target DNA; **F** 3'-labeled target DNA. Samples were analyzed on a 15% denaturing PAA gel.

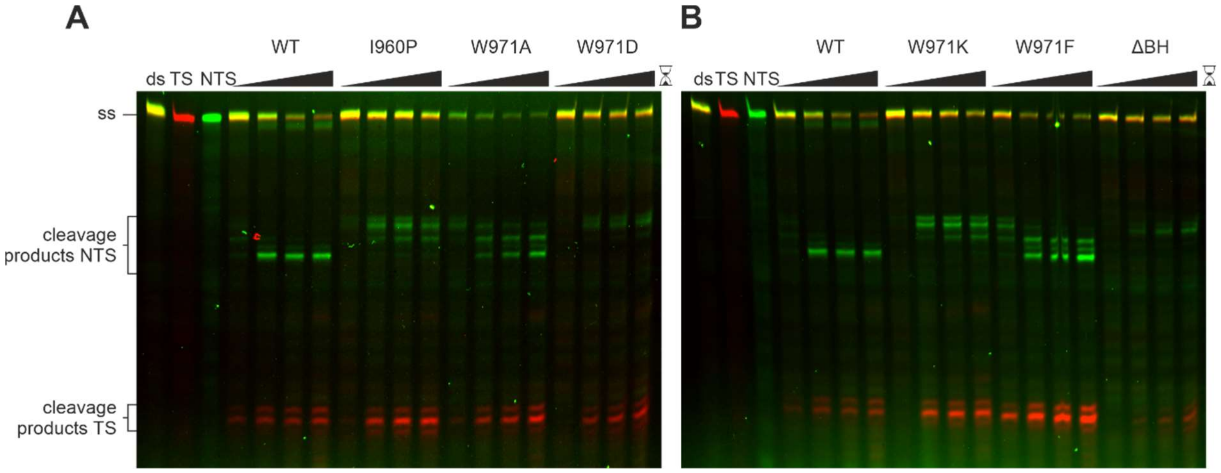


**Figure S6.** Cleavage kinetics of Cas12a mutants using a short 5' doubly labeled DNA target (target as shown in Figure S5A). The reactions contained Cas12a and crRNA (375 nM) in 7.5-fold excess over target DNA (58 nt; 50 nM). Reactions were stopped with 125 mM EDTA after different time intervals. **A** and **B** high resolution cleavage kinetics stopped after 1 min, 1 h, 2.5 h, and 5 h. Samples were analyzed on a high-resolution 15% denaturing PAA gel.

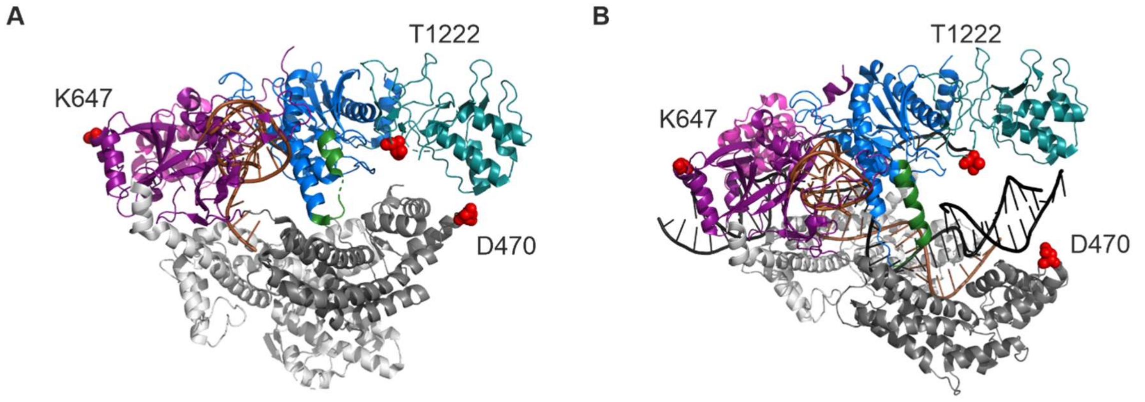


C

| | binary complex | ternary complex | ternary complex -NTS |
|---|---|---|---|
| K647-T1222 | 78.1 Å | 79.8 Å | 77.3 Å |
| D470-K647 | 98.0 Å | 104.4 Å | 105.7 Å |
| D470-T1222 | 36.0 Å | 30.8 Å | 33.2 Å |

**Figure S7. Labeling positions D470 (REC lobe), K647 (wedge domain), and T1222 (Nuc lobe) in FnCas12a.** Color coding of the Cas12a domains according to Figure 1. **A** Binary complex (PDB: 5NG6) and **B** ternary complex (PDB: 6I1K) with labeling positions shown as red spheres. **C** Distances between the $C_\alpha$-atoms of the labeling positions in binary and ternary complex.

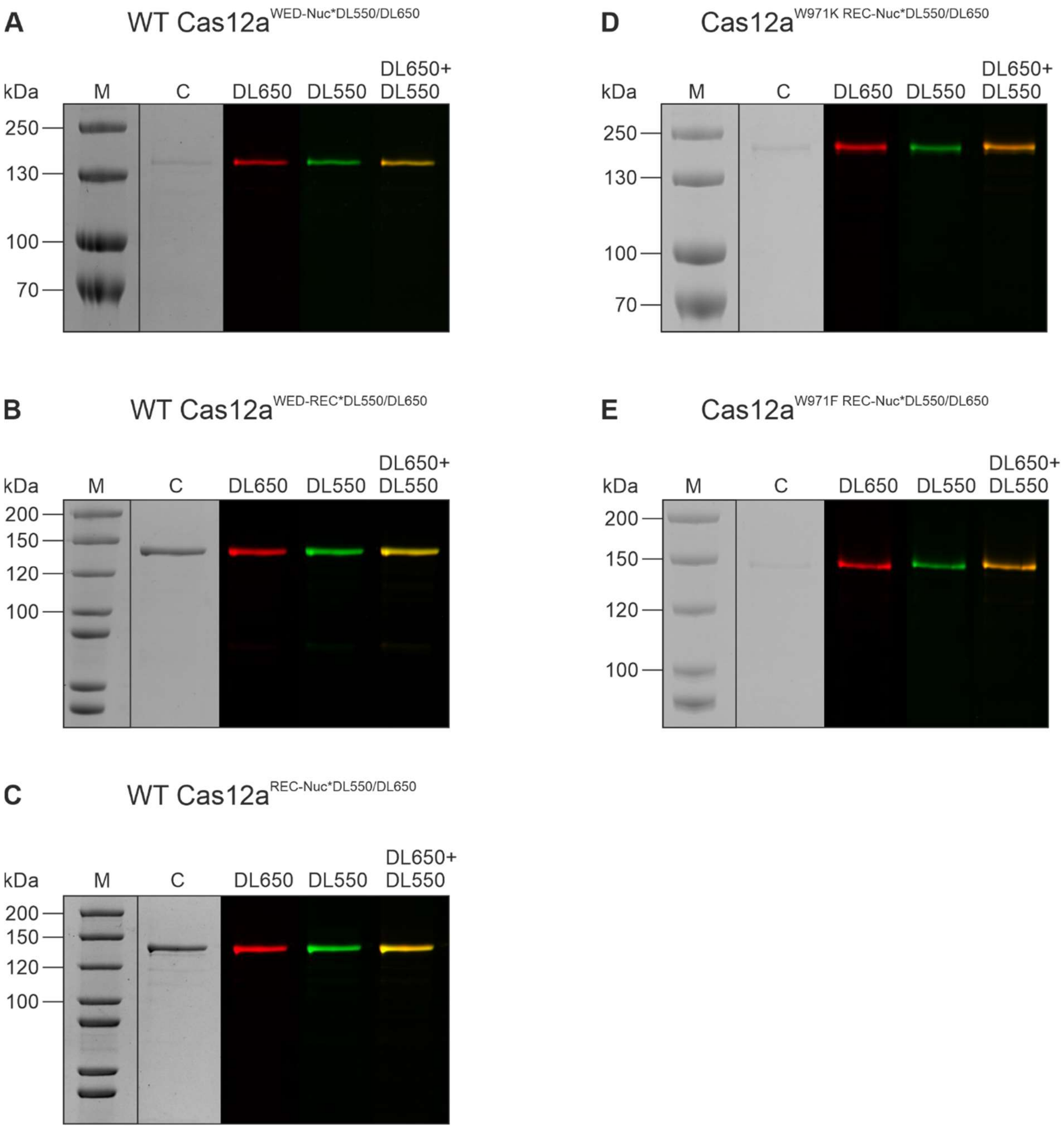


**Figure S8.** SDS-PAGE analysis of doubly labeled Cas12a variants (fluorophores are DyLight 550 (DL550) and DyLight 650 (DL650)). M: PageRuler$^{TM}$ prestained (A, D)/PageRuler$^{TM}$ unstained (B, C, E); C: Coomassie stained gel; DL650: fluorescence scan at 625-650 nm excitation for DyLight 650; DL550: fluorescence scan at 520-545 nm excitation for DyLight 550; DL650+DL550: overlay of fluorescence scans DL650 and DL550.

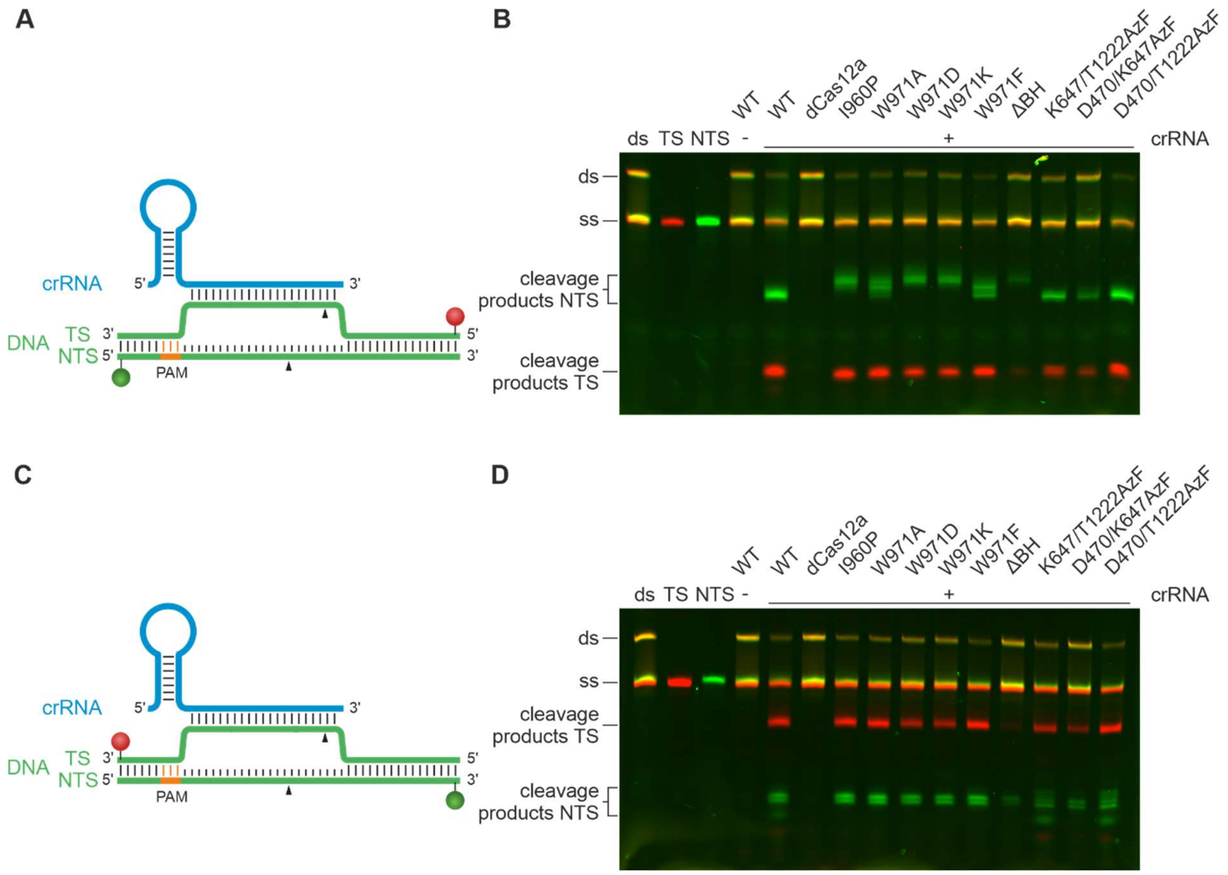


**Figure S9. Cleavage assay of Cas12a variants using a short doubly labeled DNA target.** Cleavage assay using WT Cas12a, dCas12a, Cas12a bridge helix mutants, and Cas12a with incorporated AzF. Proteins were used in a 7.5-fold excess of protein and crRNA (75 nM) over the target DNA (10 nM). Reactions were incubated for 1 h. The short target DNA (58 nt) is doubly labeled with a Cy3 label (green) at the non-target strand (nts) and a Cy5 label (red) at the target strand (ts). **A** Cy3 (green sphere) and Cy5 (red sphere) labels are coupled to the 5'-end of the DNA strands. **B** The cleaved nts shows a different cleavage pattern for the different Cas12a variants. The ts cleavage pattern is the same for all Cas12a variants. **C** Cy3 (green sphere) and Cy5 (red sphere) labels are coupled to the 3'-end of the DNA strands. **D** The cleavage patterns of target and non-target strand are the same for all Cas12a variants. Samples were analyzed on a 15% denaturing PAA gel.

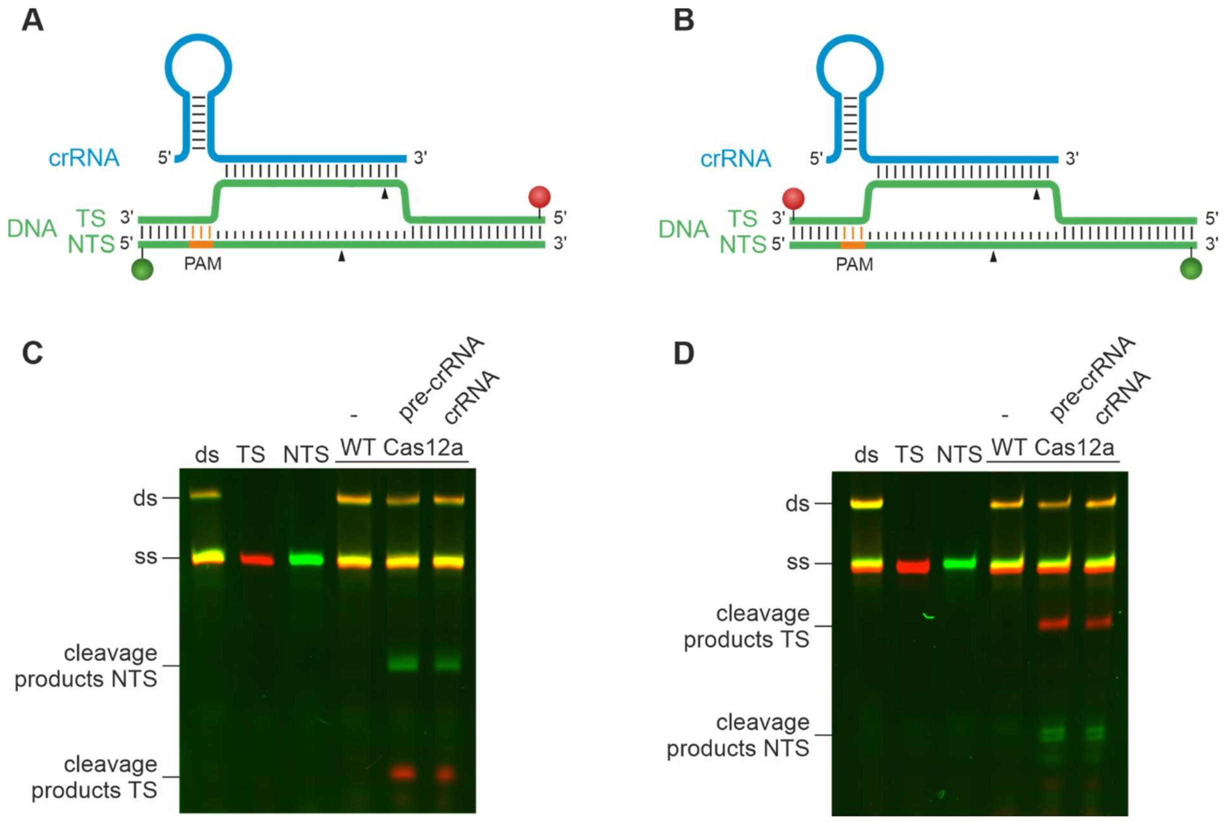


**Figure S10.** Cleavage assay using WT Cas12a and a short doubly labeled DNA target with no RNA (-), pre-crRNA, or mature crRNA. The short target DNA (58 nt) is doubly labeled with a Cy3 label (green) at the non-target strand (nts) and a Cy5 label (red) at the target strand (ts). **A** Cy3 (green sphere) and Cy5 (red sphere) labels are attached to the 5'-end of the DNA strands. **B** Cy3 (green sphere) and Cy5 (red sphere) labels are at the 3'-end of the DNA strands. **C** Cleavage assay with 5'-labeled and **D** cleavage assay containing 3'-labeled DNA (10 nM) with pre-crRNA (75 nM) or mature RNA (75 nM). Reaction products were analyzed on a 15% denaturing PAA gel. The cleavage efficiency of Cas12a (75 nM) loaded with pre-crRNA or mature crRNA is comparable.

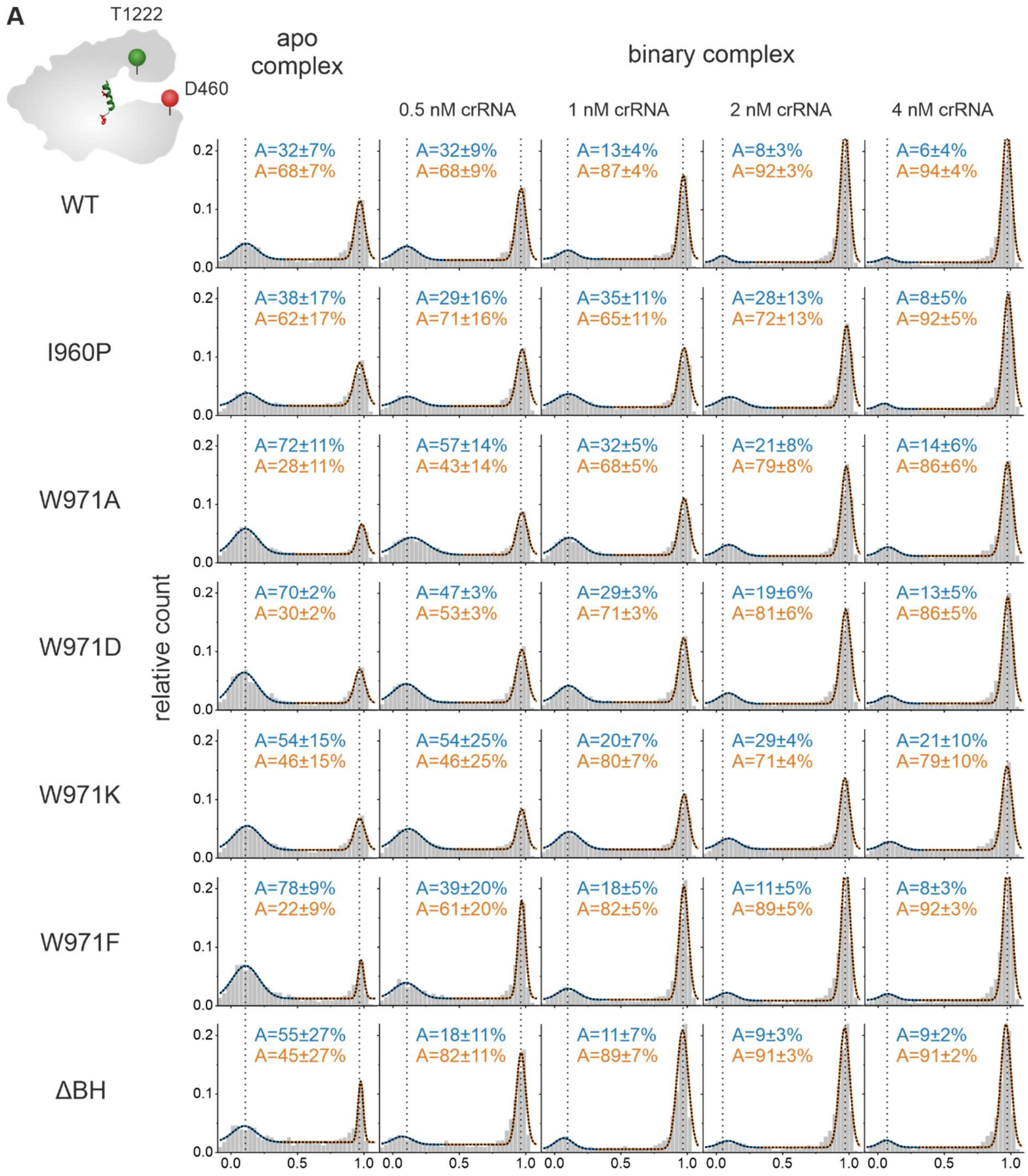
A
T1222
D460
apo complex
binary complex
0.5 nM crRNA
1 nM crRNA
2 nM crRNA
4 nM crRNA
WT
I960P
W971A
W971D
W971K
W971F
ΔBH
relative count
A=32±7%
A=68±7%
A=32±9%
A=68±9%
A=13±4%
A=87±4%
A=8±3%
A=92±3%
A=6±4%
A=94±4%
A=38±17%
A=62±17%
A=29±16%
A=71±16%
A=35±11%
A=65±11%
A=28±13%
A=72±13%
A=8±5%
A=92±5%
A=72±11%
A=28±11%
A=57±14%
A=43±14%
A=32±5%
A=68±5%
A=21±8%
A=79±8%
A=14±6%
A=86±6%
A=70±2%
A=30±2%
A=47±3%
A=53±3%
A=29±3%
A=71±3%
A=19±6%
A=81±6%
A=13±5%
A=86±5%
A=54±15%
A=46±15%
A=54±25%
A=46±25%
A=20±7%
A=80±7%
A=29±4%
A=71±4%
A=21±10%
A=79±10%
A=78±9%
A=22±9%
A=39±20%
A=61±20%
A=18±5%
A=82±5%
A=11±5%
A=89±5%
A=8±3%
A=92±3%
A=55±27%
A=45±27%
A=18±11%
A=82±11%
A=11±7%
A=89±7%
A=9±3%
A=91±3%
A=9±2%
A=91±2%
0.0
0.1
0.2
0.5
1.0

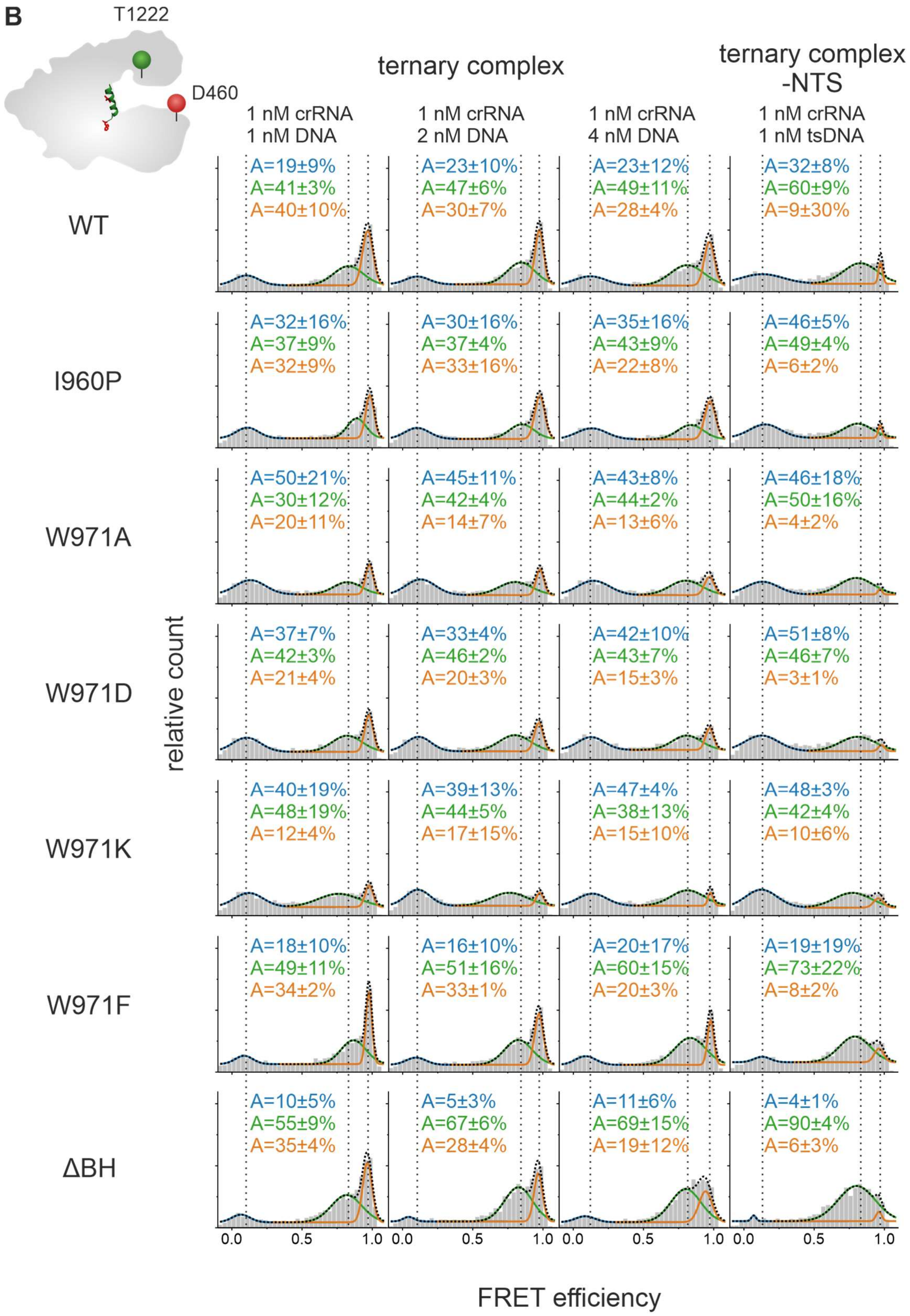


**Figure S11. FRET efficiency histograms of Cas12a$^{REC\text{-}Nuc*DL550/DL650}$ bridge helix mutants. A** Single-molecule FRET measurements of the apo complex and the binary complex with increasing crRNA concentrations: 0.5 nM, 1 nM, 2 nM, 4 nM crRNA. **B** FRET efficiency histograms of single-molecule FRET measurements of the ternary complex (1 nM crRNA; 1 nM, 2 nM, 4 nM target DNA), and the ternary complex without nts (1 nM crRNA, 1 nM ts). **A** and **B** Histograms show the data of three independent measurements. The histograms were fitted with a double or triple Gaussian function and the percentage distribution of the populations (A) are given with SEs in the histograms.

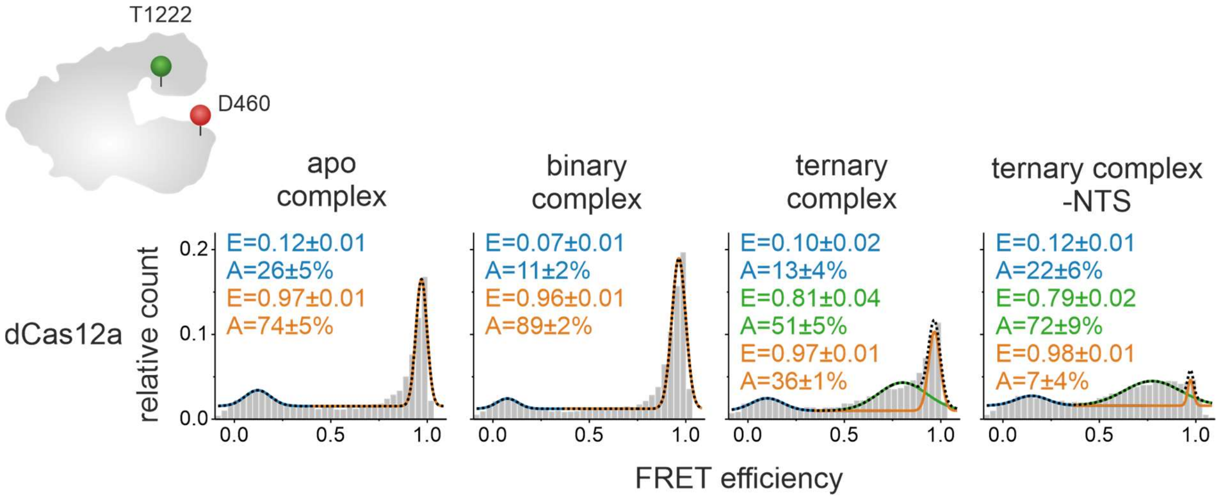


**Figure S12. FRET efficiency histograms of dCas12a$^{Rec\text{-}Nuc*DL550/DL650}$.** Single-molecule FRET measurements were conducted with the FnCas12a apo enzyme, the binary complex (1 nM crRNA), the ternary complex (1 nM crRNA, 1 nM target DNA), and the ternary complex without nts (1 nM crRNA, 1 nM ts). Histograms show the data of three independent measurements. The histograms were fitted with a double or triple Gaussian function and the mean FRET efficiencies (E) and the percentage distribution of the populations (A) are given with SEs in the histograms.

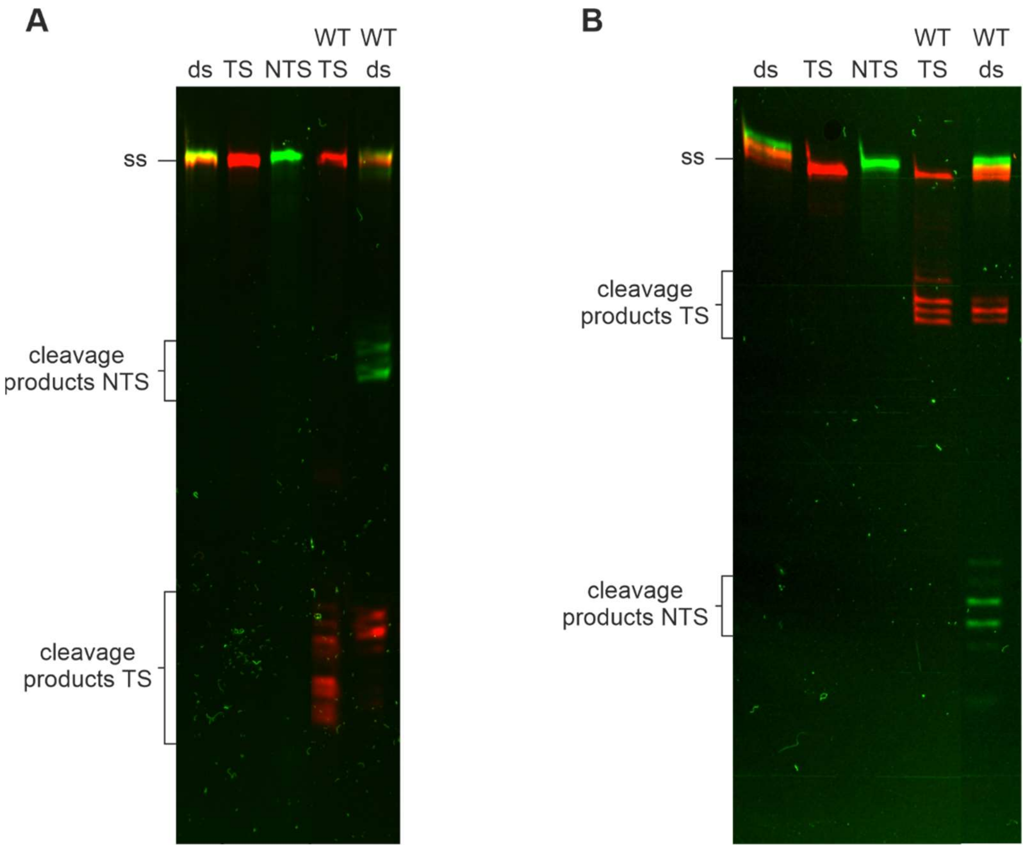


**Figure S13. Cleavage reaction containing either target strand only or a double-stranded target DNA.** The reactions contained WT Cas12a and crRNA (75 nM) in 7.5-fold excess over the target strand (ts) or ds target DNA (58 nt; 10 nM). Incubation of the reaction at 37 °C for 1 h. **A** Cleavage assay with 5'-labeled and **B** with 3'-labeled DNA strands. Samples were analyzed on a high-resolution denaturing 15% PAA gel.

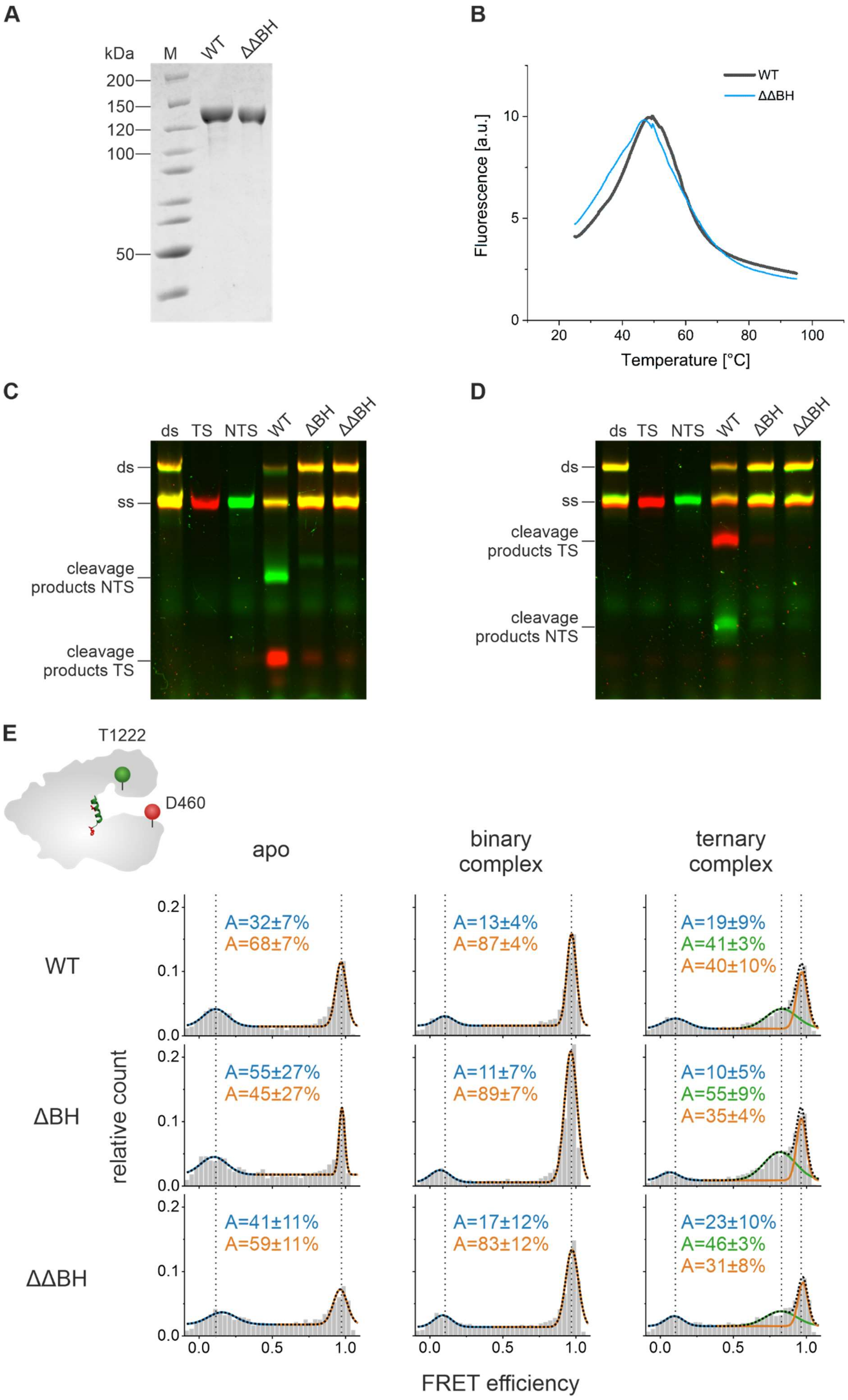

A
kDa
M
WT
ΔΔBH
200
150
120
100
50
B
Fluorescence [a.u.]
Temperature [°C]
WT
ΔΔBH
C
ds
TS
NTS
WT
ΔBH
ΔΔBH
ds
ss
cleavage products NTS
cleavage products TS
D
ds
TS
NTS
WT
ΔBH
ΔΔBH
ds
ss
cleavage products TS
cleavage products NTS
E
T1222
D460
apo
binary complex
ternary complex
WT
ΔBH
ΔΔBH
relative count
A=32±7%
A=68±7%
A=13±4%
A=87±4%
A=19±9%
A=41±3%
A=40±10%
A=55±27%
A=45±27%
A=11±7%
A=89±7%
A=10±5%
A=55±9%
A=35±4%
A=41±11%
A=59±11%
A=17±12%
A=83±12%
A=23±10%
A=46±3%
A=31±8%
FRET efficiency

**Figure S14. Characterization of Cas12a ΔΔBH mutant. A** SDS-PAGE analysis of purified WT Cas12a and Cas12a ΔΔBH (0.82 µg protein, molecular mass of FnCas12a WT: 151.9 kDa). M: PageRuler$^{TM}$ (Thermo Scientific) unstained. **B** Melting curves of WT Cas12a and Cas12a ΔΔBH. Protein Thermal Shift$^{TM}$ (Thermo Scientific) melting curves of Cas12a variants (2 µg) from 25 to 95 °C. Shown is the average of four replicates. **C** and **D** Cleavage assay of WT Cas12a and BH deletion mutants (ΔBH, ΔΔBH). A 7.5-fold excess of protein and crRNA (75 nM) over the short doubly labeled target DNA (10 nM) was used. **C** The short target DNA (58 nt) is doubly labeled with a Cy3 label (green) at the 5'-end of the non-target strand (nts) and a Cy5 label (red) at the 5'-end of the target strand (ts). **D** The short target DNA (58 nt) is doubly labeled with a Cy3 label (green) at the 3'-end of the non-target strand (nts) and a Cy5 label (red) at the 3'-end of the target strand (ts). **E** FRET efficiency histograms based on single-molecule measurements with WT Cas12a$^{REC\text{-}Nuc*DL550/DL650}$, ΔBH Cas12a$^{REC\text{-}Nuc*DL550/DL650}$, and ΔΔBH Cas12a$^{REC\text{-}Nuc*DL550/DL650}$ either as apo enzyme, the binary complex (1 nM crRNA), and the ternary complex (1 nM crRNA, 1 nM target DNA). The histograms were fitted with a double or triple Gaussian function and the percentage distribution of the populations (A) are given with SEs in the histograms.

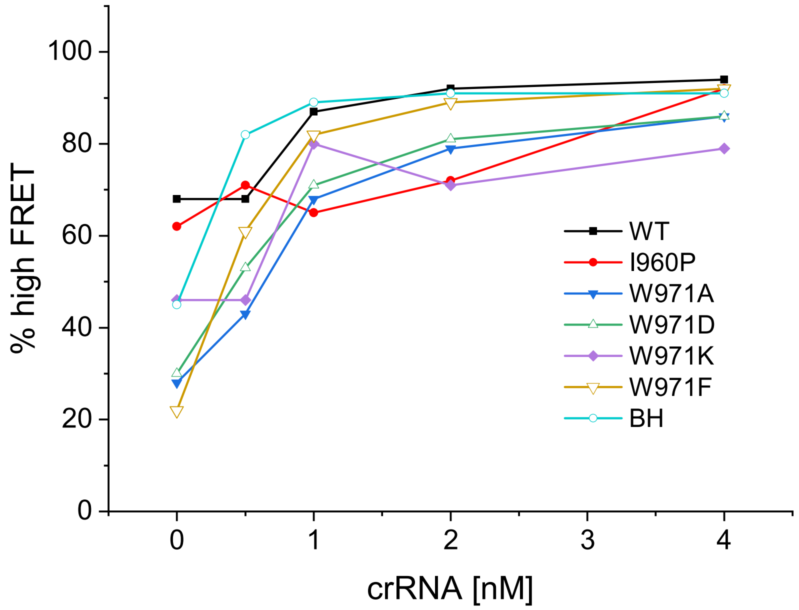


**Figure S15.** Quantification of the high FRET population in the binary complex of Cas12a$^{REC\text{-}Nuc*DL550/DL650}$ bridge helix mutants. The relative amount of the high FRET population increases upon addition of increasing concentrations of crRNA (0 nM, 0.5 nM, 1 nM, 2 nM, 4 nM). BH mutants do not adopt the conformation reflected by the high FRET population as efficiently as the wildtype enzyme.

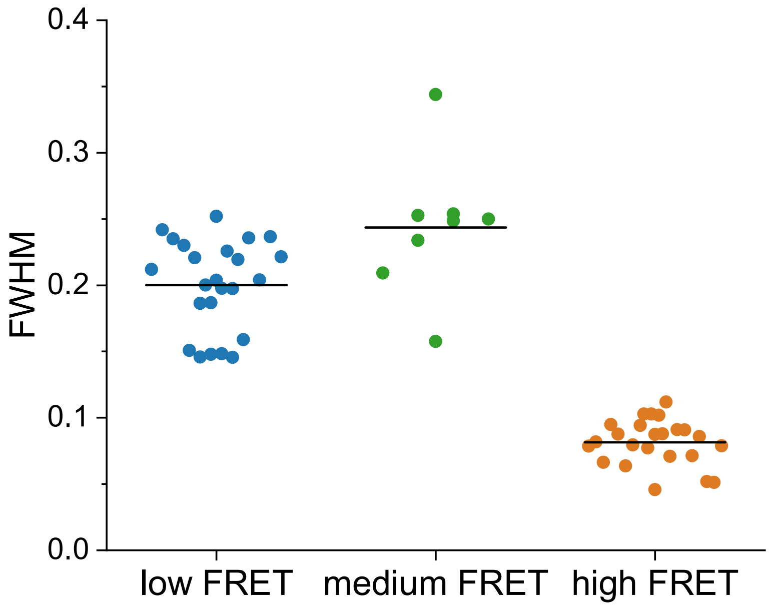


**Figure S16.** Quantification of full width half maximum (FWHM) values of WT Cas12a$^{REC\text{-}Nuc*DL550/DL650}$ and Cas12a$^{REC\text{-}Nuc*DL550/DL650}$ bridge helix mutants. Quantified are the fits displayed in figure 4 for the low, medium, and high FRET population.

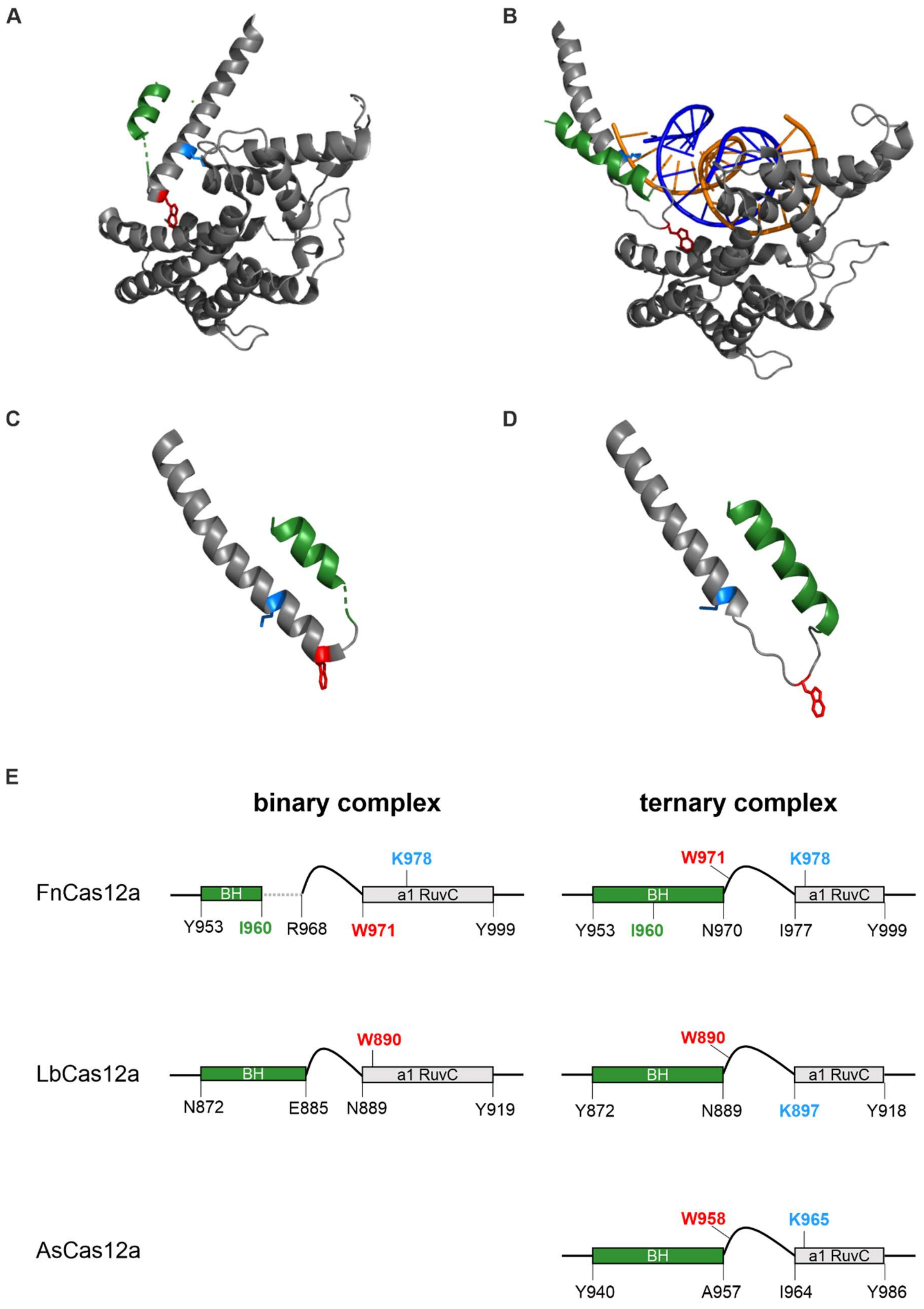


**Figure S17. Structural state of the bridge helix and helix-1 in the binary and ternary complex. A** and **B** Comparison of the structural rearrangement of the BH and helix-1 relative to the REC domain upon transition from the binary to ternary complex. **A** shows the structural elements in the binary (PDB: 5NG6) and **B** in the ternary (PDB: 6IK1) FnCas12a complex. The structures were aligned relative to the REC domain as reference point. For clarity, only the bridge helix (BH, green), helix-1 and the REC domain (dark grey) are shown. Tryptophan 971 is highlighted in red and lysine 978 in blue. **C** and **D** Comparison of the length of BH, helix-1 and connecting linker in the **C** binary (PDB: 5NG6) and **D** ternary complex (PDB: 6IK1). Tryptophan 971 is highlighted in red and lysine 978 in blue. **E** Schematic comparison of the length of the BH, the linker and the helix-1 of the RuvC domain in the binary and ternary complex. Models are based on analysis of Cas12a crystal structure from *Franciscella novicida* U112 (binary complex PDB: 5NG6, ternary complex PDB: 6I1K), *Lachnospiraceae bacterium ND2006* (binary complex PDB: 5ID6, ternary complex PDB: 5XUS) and *Acidaminococcus sp. BV3L6* (ternary complex PDB: 5KK5). The tryptophan residue that is anchored in the hydrophobic pocket is highlighted in red, the lysine residue that interacts with the DNA phosphate backbone is highlighted in blue and the isoleucine 960 residue, which was mutated to a proline in this study, is highlighted in green.

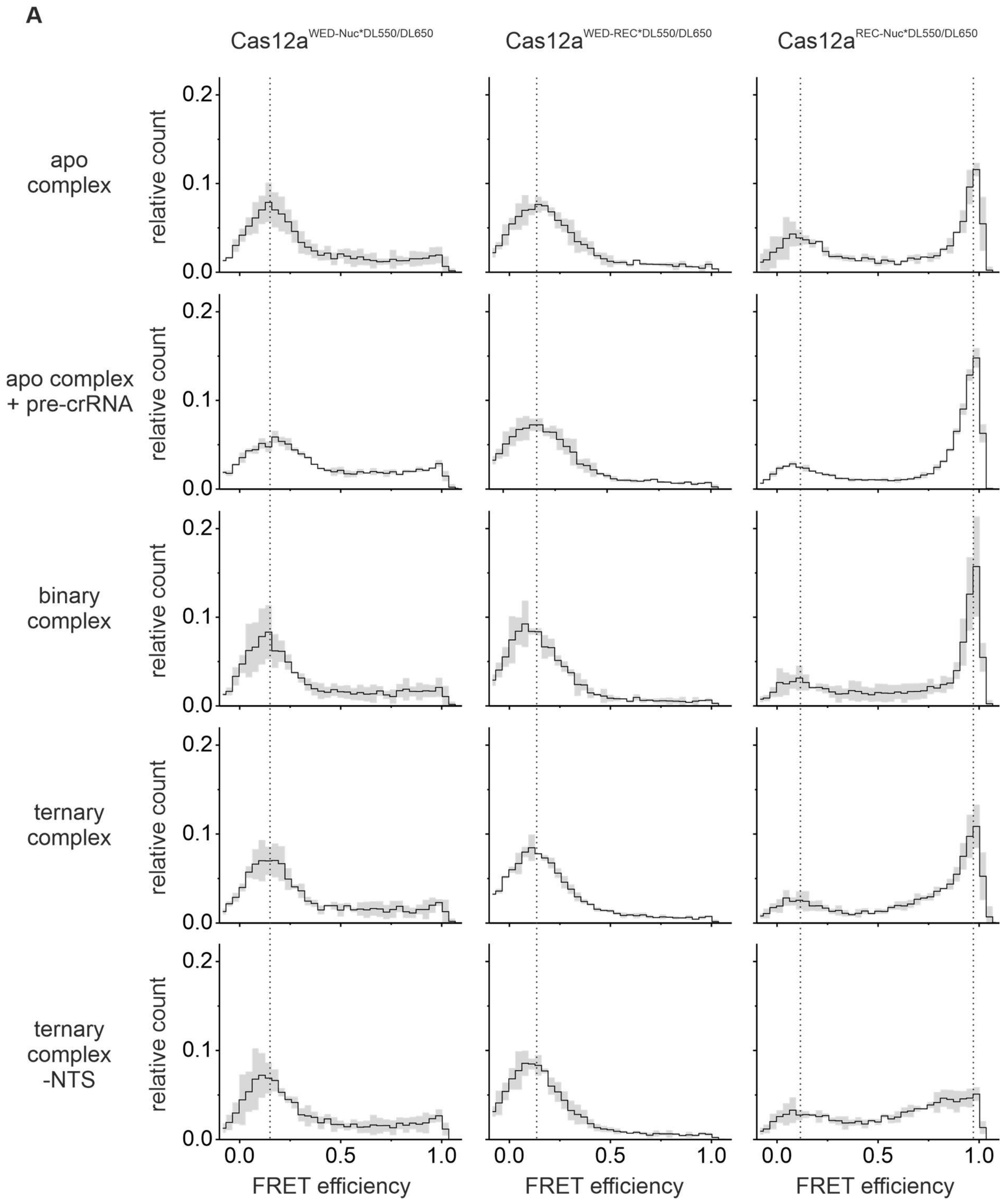

A
Cas12a WED-Nuc*DL550/DL650
Cas12a WED-REC*DL550/DL650
Cas12a REC-Nuc*DL550/DL650
apo complex
apo complex + pre-crRNA
binary complex
ternary complex
ternary complex -NTS
relative count
0.0
0.1
0.2
0.0
0.5
1.0
FRET efficiency

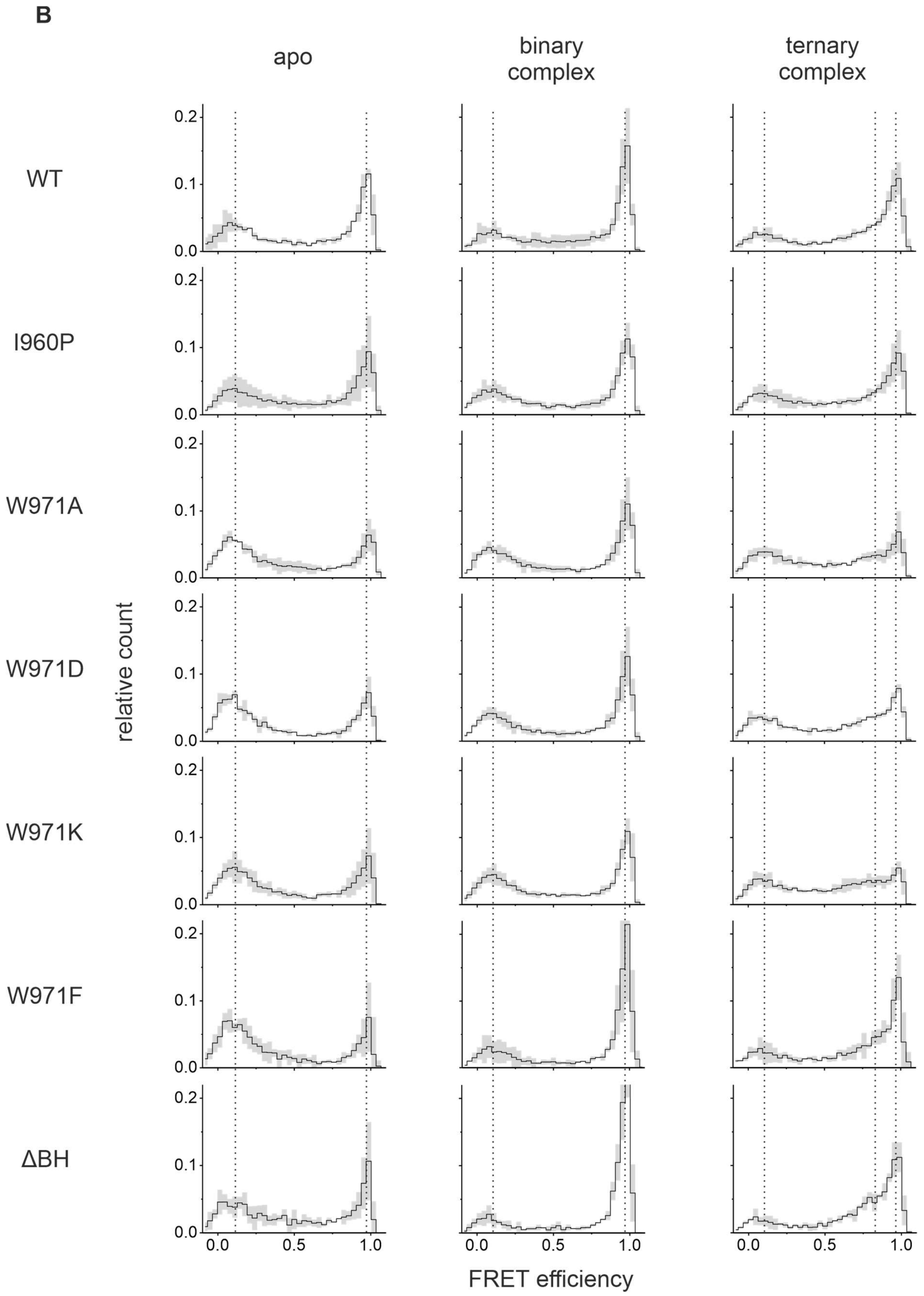


**Figure S18. Overview of all FRET efficiency histograms.** The mean FRET efficiency (black) and standard deviation of triplicates (grey) are shown. **A** FRET efficiency histograms of WT Cas12a with different labeling positions. Single-molecule FRET measurements of the apo complex, the apo complex with pre-crRNA (1 nM), the binary complex (1 nM crRNA), and the ternary complex (1 nM crRNA, 1 nM target

DNA). **B** Single-molecule FRET efficiency histograms of Cas12a$^{REC\text{-}Nuc*DL550/DL650}$ bridge helix mutants. Single-molecule FRET measurements of the apo complex, the binary complex (1 nM crRNA), and the ternary complex (1 nM crRNA, 1 nM target DNA).

Table S1. Sequences of mutagenesis primer.

| | | Sequence 5' ⟶ 3' |
|---|---|---|
| E1006A | fw | CTATTGTGGTTTTTGCGGATTTAAATTTTG |
| | rev | CAAAATTTAAATCCGCAAAAACCACAATAG |
| I960P | fw | CATGATAAGCTTGCTGCACCAGAGAAAGATAGGGATTC |
| | rev | GAATCCCTATCTTTCTCTGGTGCAGCAAGCTTATCATG |
| W971A | fw | GATTCAGCTAGGAAAGACGCCAAAAAGATAAATAACATC |
| | rev | GATGTTATTTATCTTTTTGGCGTCTTTCCTAGCTGAATC |
| W971D | fw | GGATTCAGCTAGGAAAGACGACAAAAAGATAAATAACATC |
| | rev | GATGTTATTTATCTTTTTGTCGTCTTTCCTAGCTGAATCC |
| W971K | fw | GATTCAGCTAGGAAAGACAAGAAAAAGATAAATAACATC |
| | rev | GATGTTATTTATCTTTTTCTTGTCTTTCCTAGCTGAATC |
| W971F | fw | GATTCAGCTAGGAAAGACTTTAAAAAGATAAATAACATC |
| | rev | GATGTTATTTATCTTTTTAAAGTCTTTCCTAGCTGAATC |
| ΔBH | fw | GGTAATGATAGAATGAAAACAAACGACTGGAAAAAGATAAATAAC |
| (Y953-K969) | rev | GTTATTTATCTTTTTCCAGTCGTTTGTTTTCATTCTATCATTACC |
| ΔΔBH | fw | GGTAATGATAGAATGAAAACAAACAAAAAGATAAATAACATCAAAGAG |
| (Y953-W971) | rev | CTCTTTGATGTTATTTATCTTTTTGTTTGTTTTCATTCTATCATTACC |
| D470Stop | fw | GCATAGAGATATATAGAAACAGTGTAGG |
| | rev | CCTACACTGTTTCTATATATCTCTATGC |
| K647Stop | fw | GATGATAAAGCTATCTAGGAAAATAAAGGCGAG |
| | rev | CTCGCCTTTATTTTCCTAGATAGCTTTATCATC |
| T1222Stop | fw | CAAATGCGTAACTCAAAATAGGGTACTGAGTTAGATTATC |
| | rev | GATAATCTAACTCAGTACCCTATTTTGAGTTACGCATTTG |

Table S2. FRET efficiencies of Cas12a$^{REC\text{-}Nuc*DL550/DL650}$ bridge helix mutants.

| | | **low FRET** | **medium FRET** | **high FRET** |
|---|---|---|---|---|
| **WT Cas12a** | apo | 0.12±0.04 | | 0.97±0.01 |
| | binary complex | 0.09±0.01 | | 0.97±0.01 |
| | ternary complex | 0.11±0.02 | 082±0.02 | 0.97±0.01 |
| **I960P** | apo | 0.10±0.03 | | 0.97±0.01 |
| | binary complex | 0.12±0.03 | | 0.97±0.01 |
| | ternary complex | 0.11±0.03 | 0.87±0.02 | 0.98±0.01 |
| **W971A** | apo | 0.11±0.02 | | 0.98±0.01 |
| | binary complex | 0.10±0.01 | | 0.98±0.01 |
| | ternary complex | 0.11±0.01 | 0.80±0.02 | 0.98±0.01 |
| **W971D** | apo | 0.10±0-02 | | 0.96±0.02 |
| | binary complex | 0.10±0.01 | | 0.97±0.01 |
| | ternary complex | 0.11±0.01 | 0.80±0.02 | 0.98±0.01 |
| **W971K** | apo | 0.11±0.01 | | 0.97±0.01 |
| | binary complex | 0.10±0.01 | | 0.97±0.01 |
| | ternary complex | 0.10±0.01 | 0.86±0.02 | 0.98±0.01 |
| **W971F** | apo | 0.11±0.02 | | 0.97±0.01 |
| | binary complex | 0.11±0.01 | | 0.98±0.01 |
| | ternary complex | 0.10±0.01 | 0.86±0.02 | 0.98±0.01 |
| **ΔBH** | apo | 0.10±0.05 | | 0.98±0.01 |
| | binary complex | 0.06±0.01 | | 0.97±0.01 |
| | ternary complex | 0.06±0.03 | 0.82±0.01 | 0.97±0.01 |
